# Ghrelin and risky decision-making: No credible evidence for homeostatic state modulation of neural or behavioural effects

**DOI:** 10.1101/2025.10.20.683454

**Authors:** Steven Geysen, Angela M. Brands, Heidrun Schultz, Julian Koenig, Marc Tittgemeyer, Jan Peters

## Abstract

Risk-taking is often thought to depend on homeostatic systems. However, evidence remains mixed, and the underlying mechanisms remain debated. One candidate that might drive changes in risky decision-making due to changes in homeostatic systems may be the hunger hormone ghrelin, which interacts with the dopaminergic system. In two studies, we examined the effects of experimental interventions known to affect ghrelin levels on human risky choice in healthy male participants, either after a brief fasting period (study 1, *N* = 37; *N* = 26 for fMRI analyses) or a single night of total sleep deprivation (study 2, *N* = 40; *N* = 36 for fMRI analyses).

We found no credible effects of the experimental manipulations on the proportion of risky choices. In addition, computational modelling indicated that the standard prospect theory, without accounting for choice repetition, best described the observed behaviour. However, it did not reveal consistent effects of state manipulations on model parameters, and the inclusion of manipulation-induced changes in ghrelin levels in the model likewise revealed no robust associations. FMRI analyses did not reveal effects of state manipulation on neural signatures of subjective value or choice in *a priori* defined regions of interest (bilateral ventromedial prefrontal cortex, ventral striatum, posterior cingulate cortex, and anterior cingulate cortex for subjective value; bilateral anterior insula, ventral striatum, and right medial prefrontal cortex for choice). Our results suggest that state-dependent influences on risky decision-making may be weaker than previously thought.

---

Decision-making is a fundamental neurocognitive capacity that often depends on processing the uncertainty and risk associated with available choice options (Fox & Poldrack, 2009; Johnson & Busemeyer, 2010; Morelli et al., 2022). To examine how risk is incorporated in the decision-making process, participants are typically asked to choose between options that differ in reward probability and magnitude (Lee et al., 2023; Rachlin et al., 1991). Such risky choice paradigms usually entail an optimal solution. Multiplying the reward magnitude *R* with the probability *P* yields an option’s expected value (*EV* = *R* ∗ *P*); the best option has the highest expected value. However, according to *prospect theory* (PT) by Kahneman and Tversky (1979), it is in the apparent simple calculation of expected value that people deviate from optimality. An alternative model that is frequently used to account for suboptimal risky decision-making is the *hyperbolic probability discounting model* (HPDM; Green & Myerson, 2004; Mazur, 1986). *Probability discounting* (PD) is the reduction of the subjective outcome value as a function of the probability with which it occurs (Kyonka & Schutte, 2018; Rachlin et al., 1991). While HPDM is strong in its simplicity, PT might better account for risk attitudes (Ligneul et al., 2013).

Previous research employing different paradigms to study risky decision-making have identified several involved neural systems (Cui et al., 2022; Levy, 2017; Wu et al., 2021). Risk-taking is linked to increased activity in regions associated with the neural representation of subjective reward value (Levy, 2017; Panidi et al., 2022, 2024; Seaman et al., 2018; Yao et al., 2023), including the posterior cingulate cortex (PCC), ventromedial prefrontal cortex (vmPFC), dorsolateral prefrontal cortex (dlPFC), and ventral striatum, in particular the nucleus accumbens (NAc; Bartra et al., 2013; Levy et al., 2010; Nachev et al., 2015; Pearson et al., 2011). Moreover, the anterior insula, anterior cingulate cortex (ACC), and dorsomedial prefrontal cortex (dmPFC) are important for conflict monitoring and exhibit activation increases during risk-taking (Brown & Braver, 2007; Lv et al., 2021; Von Siebenthal et al., 2020).

Several previous studies linked homeostatic state changes to the extent to which context modulates risk-taking. For example, individuals may act more risk-averse shortly after having a meal (Symmonds et al., 2010) and make riskier choices when hungry (Levy et al., 2013; Li et al., 2020; Shabat-Simon et al., 2018). Hunger and satiety, the interoceptive reports of energy deficit or surplus, respectively, are tightly controlled by well-defined neurocircuitry interactions (Beutler et al., 2017; Burnett et al., 2016; Heisler & Lam, 2017; Liu & Kanoski, 2018; Sutton Hickey & Krashes, 2020), including foremost hypothalamic areas (Siemian et al., 2021; Wallner-Liebmann et al., 2010) and the hippocampus (Davidson et al., 2007; Kanoski & Grill, 2017), amongst others, which strongly interact with the dopaminergic mesoaccumbens pathway (Cassidy & Tong, 2017; Hanßen et al., 2023; Kahn & Shohamy, 2013; Nieh et al., 2016). Hunger exerts an effect on the hypothalamic arcuate nucleus (Chen et al., 2015; Heisler & Lam, 2017; Liu et al., 2012). While hunger is operationally difficult to define, physiological proxies linking hunger state with metabolic signals may be relevant in informing state-adaptive processes. To that end we considered ghrelin as a critical peptidergic signal of state mediating hunger-driven behavioural phenotype (Müller et al., 2015; Zanchi et al., 2017). Ghrelin levels increase in anticipation of food intake, and decrease following consumption (Cummings et al., 2001; Deschaine & Leggio, 2022). Ghrelin is mainly produced in the stomach (Kojima et al., 1999; Tschöp et al., 2000) and regulates feeding behaviour, metabolism, and memory through the vagus nerve (Date et al., 2006; Wren et al., 2000; Yanagi et al., 2018). Hunger elicits approach and motivation for food-seeking behaviour. Ghrelin is again a key component here since these behaviours are found to be mediated by ghrelin-induced dopaminergic activity in the VTA and NAc (Abizaid et al., 2006; Al Massadi et al., 2019; Schulz et al., 2023; Stievenard et al., 2017). At the same time, increased dopamine (DA) levels may be linked to increases in risk-taking (Rigoli et al., 2016; Rutledge et al., 2015), although the exact nature of this relationship is still unclear (Castrellon et al., 2019). The suspected role of ghrelin in risky choice behaviour is supported by the observation that participants with higher ghrelin levels scored higher on reward sensitivity and lower on punishment sensitivity (Ralevski et al., 2018), which may be conceptually related to risk-taking. In addition, gamblers continue to gamble longer in the face of losses when their ghrelin levels are higher (Sztainert et al., 2018).

Insulin is another peptide affected by feeding behaviour with the ability to interact with the dopaminergic reward system via the VTA and NAc (Liu & Borgland, 2019; Thanarajah et al., 2019).

Along similar lines, sleep and circadian rhythms affect cognition (see e.g., Harrison & Horne, 2000; Krause et al., 2017; Lowe et al., 2017). Sleep deprivation shows effects on risky choice behaviour similar to those of hunger, with some studies showing that sleep deprivation is associated with increased risk-taking (Brunet et al., 2020; Dickinson et al., 2022; Salfi et al., 2020; Venkatraman et al., 2007; Womack et al., 2013). However, other studies show no effect at all (Mao et al., 2023; Maric et al., 2017; Mullette-Gillman et al., 2015). Sleep deprivation may decrease activity in the thalamus and vmPFC (Menz et al., 2012; Thomas et al., 2000; Venkatraman et al., 2011). Moreover, sleep deprivation may increase general reward processing in the mesolimbic system (which includes the VTA and ventral striatum), decreasing the ability of discriminating between different rewards (Krause et al., 2017).

Similar to the effects of hunger discussed above, effects of sleep deprivation may in part result from peptidergic changes (Kim et al., 2015). For example, adenosine interacts with DA (Basheer et al., 2004; Fuxe et al., 2010), and levels increase following sleep deprivation (Basheer et al., 2004; Elmenhorst et al., 2007). Importantly, sleep deprivation also affects several peptides involved in metabolic functions (Kulkarni et al., 2024; van Cauter et al., 2007). Ghrelin plays a role in the modulation of sleep (Deschaine & Leggio, 2022; Hirotsu et al., 2015), and increased ghrelin levels are linked to the acute effects of sleep deprivation (AlDabal, 2011; Schmid et al., 2008; Taheri et al., 2004), although this association may be weaker than the association between hunger and ghrelin (Soltanieh et al., 2021). Leptin is another peptide that is affected by both metabolism (Austin & Marks, 2009; Klok et al., 2007) and restricted sleep duration (Spiegel, Leproult, et al., 2004; Spiegel, Tasali, et al., 2004; van Egmond et al., 2023). Leptin signals energy levels over longer time periods (Ahima & Flier, 2000; Friedman, 2019) and it does not seem to depend significantly on individual meals (Austin & Marks, 2009; Friedman, 2019; Korbonits et al., 1997). Interestingly, leptin does interact with the VTA (Fulton et al., 2006; Geisler & Hayes, 2023; Opland et al., 2010), and higher leptin levels may correlate with more risk-taking behaviour (Chang et al., 2016; Symmonds et al., 2010).

In addition to potential state-dependencies, decision-making is susceptible to a host of potential biases and choice policies. Perseveration, the tendency to repeat previous actions, is a prominent example. Perseveration is defined as a reward-independent process, which depends only on a participant’s choice history (Bornstein & Banavar, 2023; Miller et al., 2019). This could be based on information from the previous trial only (*first-order perseveration*; FOP) or information spanning a longer period (*higher-order perseveration*; HOP). Several studies have shown that behaviour is better described by models with, rather than without, a perseveration term (Palminteri, 2023; Wiehler et al., 2021). We therefore extended the risky choice models with perseveration terms (see Methods) and tested whether such models outperformed their respective traditional versions.

We investigated risky decision-making under two state manipulations. Participants performed a risky choice task (Peters & Büchel, 2009) during functional magnetic resonance imaging (fMRI) in two separate within-subjects studies. In study 1, participants performed the task hungry versus sated. In study 2, a different group of participants performed the task once following a single night of total sleep deprivation, and once following normal sleep. Behaviour was examined using model-agnostic measures as well as a detailed process-level analysis via hierarchical Bayesian modelling.

Based on previous research (Dickinson et al., 2022; Levy et al., 2013; Pietrzak et al., 2023; Shabat-Simon et al., 2018; Venkatraman et al., 2007, 2011) and considerations outlined above, we predicted participants in both the hungry and sleep-deprived conditions to show more risk-seeking behaviour compared to the sated and normal sleep conditions, respectively. In addition, we investigated if such potential manipulation effects could be explained by manipulation effects on individual-participant ghrelin levels.

## Methods

We report on two separate but related studies, examining the effect of state modulation on human risk-taking. The data collected for study 1 (hunger) have previously not been analysed or published. The data collected for study 2 (sleep deprivation) were part of a previous study for which data from another task have previously been published (Rihm et al., 2019). In the current study, we focused on the data of the probability discounting tasks included in both studies, which have previously not been analysed and published. We preregistered our analyses (https://osf.io/nk7v4) after the data collection.

### Participants

In study 1, 37 healthy right-handed men, ranging in age from 19 years to 36 years old (*M* ± *SD* = 25.62 ± 3.51), volunteered. The behavioural analysis was done for all 37 participants, the neuroimaging analysis contained the data of 26 participants (*M* = 25.35 ± 2.76, range = 19 − 30). Seven participants did not have complete fMRI data due to scanning problems. One participant was excluded due to peculiar choice behaviour, making it impossible for the first-level choice model to fit to their imaging data. An additional three participants were excluded due to problems with the analysis of the blood samples.

In study 2, 40 healthy right-handed men volunteered. Their age ranged between 19 years and 33 years old (*M* = 25.68 ± 3.66). The behavioural analysis was done for all 40 participants, the neuroimaging analysis contained the data of 34 participants (*M* = 26.12 ± 3.59, range = 19 − 33). Two participants did not have complete fMRI data due to scanner problems. Three participants were excluded due to missing endocrine data. One additional participant was excluded due to a BMI value > 25 kg/m².

In both studies, participants performed a battery of different tasks (including the PD task) in two sessions, once in a hungry (study 1) or sleep-deprived state (study 2) and once in a sated state or after a normal sleep, respectively, as a baseline measure. During both sessions, functional MRI was recorded during task performance. Prior to participation, all participants provided written and informed consent, and all study procedures were approved by the local Institutional Review Board (Ethics Committee of the Hamburg Board of Physicians, codes PV3697 and PV4377).

### Experimental Design

Both studies consisted of three sessions, each scheduled on a different day: one behavioural pretest and two counterbalanced experimental sessions. During the pretest, which took place two to five days before the first experimental session, participants completed a risky choice task for the calculation of each participant’s *indifference point*, i.e., the amount for which participants were indifferent between the reference option and the risky option. This was done by adjusting the amount of the risky option after two successive choices for the same option (reduced after two successive risky choices, increased after two successive safe choices). Indifference points were calculated for each reward probability, and used to compute participant-specific trials for the fMRI sessions (see below).

The experimental sessions were separated by one week. During the experimental sessions, right before participants entered the MRI scanner to start the task, blood samples were collected to analyse participants’ ghrelin, leptin, cortisol, insulin, and glucose levels.

### Study 1

This study had a within-subjects design where participants were either sated or hungry in the experimental sessions. Conditions were counterbalanced across participants. On the day of testing the hungry condition, participants were instructed to have their usual breakfast before 9: 00 a.m., skip lunch, and not consume any snacks or caloric drinks. The average time since their last meal was 8 hours and 19 (±27) minutes. In the sated condition, participants were instructed to not deviate from their standard eating behaviour. The average time since the last meal in the sated condition was 2 hours and 52 minutes (±1 hour 8 minutes). On experimental days, participants arrived at the institute at 4: 00 p.m. They rated their hunger level (1 = not at all hungry, 7 = very hungry) on a visual analogue scale and received instructions on the risky choice task.

### Study 2

This study had a within-subjects design where participants were either well rested or sleep deprived in the experimental sessions. Conditions were counterbalanced across participants. Participants were initially told they could be in the same condition twice. This was done to prevent participants from preparing for the upcoming condition by sleeping more during the day. The evening before each experimental session, at 8: 00 p.m., participants came to the institute and received a standardised meal. For the sleep deprivation condition, participants stayed at the institute and were kept awake for one whole night. In the normal sleep condition, participants were sent back home with an Actiwatch 2 (Philips Respironics) and the instruction to sleep normally. The average sleep duration was 6 hours and 44 (±56) minutes. In both conditions, participants were asked to not consume any food or caloric drinks. The data was collected the following morning between 7: 30 a.m. and 9: 30 a.m. For more details on the experimental design of study 2, see Rihm and colleagues (2019).

### Risky Choice Task

In both studies, participants performed a variant of the probability discounting task described by Peters and Büchel (2009). On each trial, participants were presented with a hypothetical monetary reward and the probability with which it may be received. The reward magnitude (*R*) was sampled from a Gaussian distribution with the indifference point of the participant as mean, and ranged between €20.50 and €80. The magnitude was presented alongside one of six different reward probabilities (*P* ∈ {0.17, 0.28, 0.54, 0.84, 0.96, 0.99}). This risky option was compared to the safe reference option, which was fixed to €20 guaranteed for the duration of the task. The safe option was not presented on the screen. Participants used an MRI-compatible response box to indicate which of the presented options they preferred.

The average duration of each trial was 14 seconds. A new trial was indicated with a green dot, shown for 500 *ms*. Next, we presented the risky option for 2,500 *ms*, followed by a red dot (between three and seven seconds, drawn from a uniform distribution). Then, a red cross (safe option) and green checkmark (risky option) appeared on either side of the screen. A box appeared around the selected option for two seconds. At the end of each trial, another red dot was presented for three to seven seconds. See Figure 1 for an example trial.

**Figure 1.**
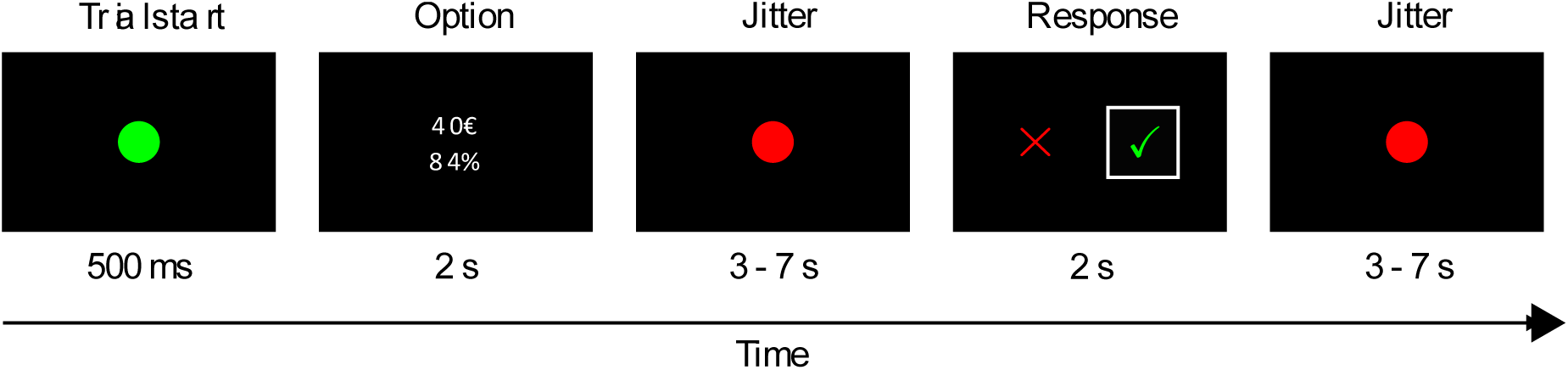
Schematic Illustration of Example Trial. Participants (*N*_1_ = 37, *N*_2_ = 40) on each trial (*T*_1_ = 96, *T*_2_ = 48) chose between a safe option (€20 with 100%) and a risky option presented on the screen. The reward magnitude for the risky option ranged between €20.50 and €80. Reward probabilities ranged between 17% and 99% (see methods section). Adapted from Peters and Büchel (2009).

The two studies differed in the number of trials per condition. In study 1, participants completed 96 trials per condition (total duration ≈ 22 minutes). In study 2, participants completed 48 trials per condition (total duration ≈ 11 minutes).

### Blood sampling

Blood samples were collected in BD P800 tubes (BD Biosciences, Heidelberg, Germany) filled with a K2EDTA anticoagulant for ghrelin and glucose plasma concentration determination and in Sarstedt Serum Gel Monovettes (Sarstedt, Nümbrecht, Germany) for leptin, cortisol, and insulin serum concentration determination. Plasma samples were immediately centrifuged for 10 min at 1.2 g at 4°C. The supernatant was immediately pipetted off and stored at −80°C. Serum samples soaked for at least 45 min in the gel solution, and were centrifuged for 10 min at 2.0 g at room temperature, and the supernatant was pipetted off and stored at −80°C. All hormone analyses were conducted by the LADR laboratory in Geesthacht, Germany. De-acylated ghrelin was analysed with an immunoassay from Bertin Pharma (Paris, France), leptin with a sandwich enzyme-linked immunosorbent assay (ELISA) from DRG (Marburg, Germany), cortisol and insulin with an electro-chemiluminescence iImmunoassay (ECLIA) method from Roche (Basel, Switzerland), and glucose with a photometric AU 5800 from Beckmann Coulter (Krefeld, Germany). For more details, see Rihm and colleagues (2019). We used de-acylated ghrelin levels (referred to as ghrelin) for the analyses.

### Functional MRI Data Acquisition

Neuroimaging data were obtained using a Siemens Magnetom Trio 3*T* whole-body scanner with a 32-channel head coil. For the functional images, we used single-shot echoplanar imaging and simultaneous multiband acquisitions with *TR* = 2,260 *ms*, *TE* = 30 *ms*, number of slices = 60, flip angle = 80°, and voxel size = 1.5 ∗ 1.5 ∗ 1.5 *mm*.

### Behavioural Data Analysis

The proportion of risky choices was used as a model-agnostic index for risk-taking propensity. For each study we used Bayesian paired samples t-tests to analyse the evidence in favour of increased risky choices in the experimental conditions. The proportion of risky choices was then further examined depending on the reward probability with a Bayesian ANOVA. The reported Bayes factors (*BF*) were interpreted following the guidelines proposed by Jeffreys (Jeffreys, 1998; Lee & Wagenmakers, 2013).

### Computational Modelling

To quantify potential effects on latent subcomponents of risk-taking, we set up and compared several cognitive computational models. These are described in the following section and differed in terms of the valuation component (e.g., prospect theory versus hyperbolic probability discounting) and the choice component describing how these subjective values guide participants’ decisions (e.g., models with first-order-perseveration (FOP) versus higher-order-perseveration (HOP) versus no perseveration). See Table 1 for an overview of the model space.

**Table 1.** Overview Model Space. The different models and their free parameters that were compared during the analyses. The underlined model provided the best fit to the behavioural data, based on the WAIC scores (see Figure S4 in the Supplementary material for the model comparison).

| Model | Free parameters | Equation |
| --- | --- | --- |
| Baseline model |  |  |
| First-order perseveration (FOP) | $\beta$ | Equation 6 |
| Higher-order perseveration (HOP) | $\alpha, \beta$ | Equation 6 |
| <u>Prospect theory model</u> | $\delta, \gamma, \beta$ | Equation 2 |
| Prospect theory with FOP | $\delta, \gamma, \beta$ | Equation 2, 7 |
| Prospect theory with HOP | $\delta, \gamma, \alpha, \beta$ | Equation 2, 7 |
| Hyperbolic probability discounting model | $k, \beta$ | Equation 4 |
| Hyperbolic discounting model with FOP | $k, \beta$ | Equation 4 |
| Hyperbolic discounting model with HOP | $k, \alpha, \beta$ | Equation 4, 7 |

### Prospect Theory Model

The PT model posits that the subjective value *SV*_*t*_ of the risky option at trial *t* is calculated by multiplying the reward magnitude *R*_*t*_ with the subjective weight of the reward probability *w*(*P*_*t*_):

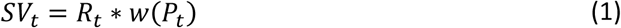

The subjective weight of the reward probability is calculated as

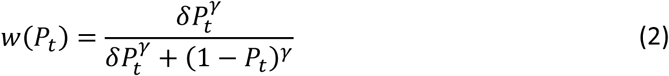

Here, the free parameter δ represents the attractiveness of risk and indicates the height of the weighting function. The sensitivity to probabilities γ indicates the degree of curvature of the probability weighting function (Goldstein & Einhorn, 1987; Kahneman & Tversky, 1979; Lattimore et al., 1992; Ligneul et al., 2013; Tymula et al., 2023).

The subjective value and the reward magnitude of the safe option (€20) are then used to estimate the probability of selecting the risky option following the Bernoulli distribution *p_risky_* ∼ *Bernoulli*_*t*_)) where the standard deviation σ_*t*_ is derived from

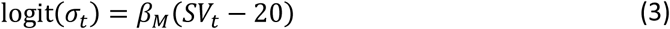

with higher β_*M*_ indicating higher stochasticity (i.e., random exploration), and lower values depict value-focused behaviour (i.e., more exploitation).

All models include the Bernoulli distribution to estimate the probability of selecting the risky choice option based on the resulting subjective values.

### Hyperbolic Probability Discounting Model

The hyperbolic probability discounting model (Green & Myerson, 2004; Johnson et al., 2020; Kyonka & Schutte, 2018; Mazur, 1986; Rachlin et al., 1991) calculates *SV*_*t*_ of *R*_*t*_ with

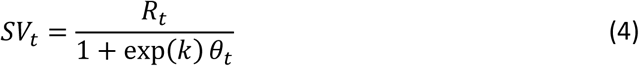

where θ_*t*_ is the odds against winning the gamble of the risky option and is derived according to

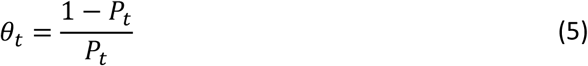

*P*_*t*_ is the reward probability. The discount rate *k* (modelled here in log-space) quantifies how rapidly subjective values change as a function of probability and thus represents the participant’s risk preference. Greater values of *k* reflect a steeper discounting of value over probability, and therefore greater risk aversion.

### Prospect Theory Model with Perseveration

Since we expected that accounting for choice repetition could further improve model fit, we added a HOP term, as proposed by Miller and colleagues (2019), to the PT model (Equations 1 and 2). This HOP term tracks the choice history in a matrix ***H*** of size *O* ∗ *T* (number of options ∗ number of trials). The matrix was initialised with 0 and is updated at each trial, according to

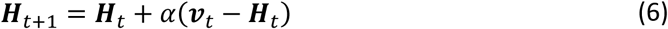

with α as step-size parameter depicting the extent to which previous actions further in the past are updated, and ***v***_*t*_ is a contrast vector, of which all elements are 0 except for the one corresponding to the selected option at trial *t*.

To estimate the probability of selecting the risky option, σ_*t*_ of the Bernoulli distribution included the weighted HOP matrix, resulting in

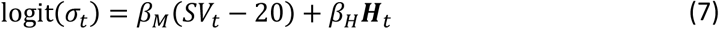

where scaling parameters β_*M*_ and β_*H*_ control the weight of the model and perseveration term, respectively.

### Hyperbolic Probability Discounting Model with Perseveration

The same principle is applied to the HPDM. The perseveration term is the same as in Equation 6. The only difference is the calculation of *SV*_*t*_ in Equation 7 as it is following the hyperbolic probability discounting model (Equation 4).

For both models with a perseveration term, we had both FOP and HOP versions. The FOP term is a restricted case of HOP, where the step-size parameter α is fixed to 1 instead of being able to vary between 0 and 1. In other words, the FOP term took only the previous trial into account for the calculation of the repetition strength, while the HOP term used all previous trials for the calculations.

### Baseline Models

The models with a value function were compared to two baseline models, consisting of only a perseveration term. This was either the FOP term or the HOP term.

### Model Fit and Comparison

The different models were fitted using hierarchical Bayesian parameter estimation in RStan (version 2.26.22; Stan Development Team, 2023). In a first step, all models were fitted to data from each condition separately, resulting in separate group-level distributions per condition and subject-level parameter. Prior distributions for group-level means were set to be normally distributed (μ = 0, σ = 5), truncated at 0. To determine the best-fitting model, we compared their widely applicable information scores (WAIC; Watanabe, 2010). Here, lower scores indicate a superior fit. For each condition, in both studies, the PT model without perseveration term had the lowest WAIC score and was therefore regarded as best fitting model to the data (see Figure S4 in the Supplementary material).

In a second step, we fitted data from both conditions per study using an adapted version of the PT model (Equation 2), to directly account for condition effects (*shifts*). In order to estimate these effects, we adapted the model so that, for each free parameter, an additional parameter was added that accounts for parameter changes due to the experimental manipulation. In case of the subjective probability weight this results in

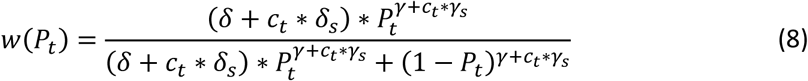

Here, *c*_*t*_ depicts a binary indicator which equals zero for the baseline condition and 1 otherwise. This way, the shift parameters (e.g., δ_*s*_ and γ_*s*_) account for the change in parameter estimates due to the condition. For each model, we ran four chains with 2,000 samples, half of which were discarded as warmup.

To examine the effects of fasting and sleep deprivation on the different parameters, we focused on the posterior distributions of the group-level parameter derived from the best fitting model. To this end, we provided the 85% and 95% highest density intervals (HDI) and *dBF*_10_ from Bayesian paired samples t-tests to quantify such potential effects.

### Ghrelin Analysis

We used Bayesian paired samples t-tests to analyse the evidence in favour of increased ghrelin levels in the experimental conditions (hungry or sleep deprived) compared to the baseline conditions. Similarly, the effect of ghrelin on the proportion of risky choices was analysed by comparing a Bayesian linear model with ghrelin and condition as predictors of the proportion of risky choices, to a model with only condition as predictor. Both models included a random intercept for the participants.

To analyse the effects of ghrelin on the parameters of the PT model, we allowed each model parameter to vary according to the difference in ghrelin levels between experimental conditions. For this we added a ghrelin compound value to each parameter which accounts for the difference in ghrelin levels between the conditions. Using δ as example, its ghrelin compound value δ_*g*_ is calculated as

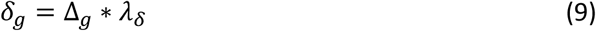

δ_*g*_ contains the difference in ghrelin between conditions Δ_*g*_, and the ghrelin coefficient λ_δ_ for the model’s parameter, in this example δ. The same is done for γ_*g*_ and β_*g*_. This resulted in the final PT model where the subjective probability weight (Equation 2) is

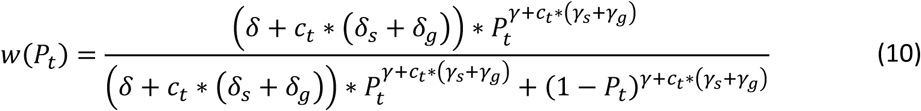

### Neuroimaging Analysis

Preprocessing and analyses of neuroimaging data were done with SPM12 (Penny et al., 2006), following the same procedure as Rihm and colleagues (2019). The first five scans were removed, the remaining images were realigned to the first scan and unwarped to correct for motion. As Rihm and colleagues, we too skip slice-timing correction. The anatomical scans were coregistered to the mean EPI image and segmented in grey and white matter, and cerebrospinal fluid (CSF). For each participant, and each condition, we computed noise regressors by applying principal components analysis (PCA) on a mask image of both white matter and CSF.

Imaging analysis focused on two effects. First, we examined neural correlates of subjective value effects followed by an investigation of the effects related to risky versus safe choices.

### Subjective Value Effects

The examination of neural correlates of subjective value effects consisted of replication analyses and analyses of the effect of the state conditions.

For the first-level (within-subject) analyses, we used a different general linear model (GLM) in native space for each participant, for each condition separately. These GLMs included a stick function for the risky option onset, the participant’s z-scored subjective value as computed by the best fitting model as parametric regressor, and the squared z-scored subjective value as a second parametric regressor. The PT parameter values required to calculate the subjective values were estimated by fitting the winning model to the subset of participants used in the fMRI analysis and taking the mean values of each participant’s individual estimated parameter distribution. The GLMs additionally contained a regressor modelling error trials (e.g., trials without responses), as well as noise regressors from the PCA as covariates. All regressors in the model, except for the noise regressors, were convolved with the canonical hemodynamic response function. We applied the default high-pass filter (HPF) cut-off of 128 seconds. Contrast images were created for the linear effects of subjective value. We normalised the segmented anatomical images and the contrast images to a standard space coordinates, with a voxel size 1.5 ∗ 1.5 ∗ 1.5 *mm* (IXI549Space of SPM12, similar to Montreal Neurological Institute (MNI) space). The contrast images were smoothed with a 6 *mm* full-width at half-maximum (FWHM) isotropic Gaussian kernel.

During the second-level analyses, we investigated the main effect of subjective value by comparing the mean BOLD responses of both conditions (experimental and baseline condition) to zero using a t-test. We then tested the effects of state-dependencies on the aforementioned value effects by comparing the mean BOLD responses of each experimental condition to the mean BOLD responses in the baseline conditions using a paired t-test. For both tests, we corrected for multiple comparisons by using small-volume corrections (*p* < .05, familywise error (FWE) corrected) in regions related to value processing. Therefore, we used the region of interest (ROI) mask provided by the Rangel Neuroeconomics Laboratory (https://www.rnl.caltech.edu/resources/index.html). This mask is based on two meta-analyses (Bartra et al., 2013; Clithero & Rangel, 2014) and defines ROIs which correlated positively with reward value (bilateral vmPFC, ventral striatum, PCC, and ACC).

To investigate the effect of ghrelin in each study, we used again two GLMs, one for each study, to correlate the z-scored difference in ghrelin levels between conditions (baseline versus manipulation) with the parametric modulated brain activation. We used the meta-analysis of Schulz and colleagues (2023; https://identifiers.org/neurovault.collection:13247) as mask for small-volume correction. This mask includes the insula, ventral striatum, and hypothalamus amongst others.

### Choice Effects

To examine effects related to safe versus risky choices, replication analyses and analyses of the effect of the metabolic state conditions were performed. For the first-level analysis in native space, the GLMs (one for each study) included stick functions for the risky option onsets of safe choices and risky choices, an regressor modelling error trials for risky onsets, and noise regressors from the PCA as covariates. All but the noise regressors in the model were convolved with the canonical hemodynamic response function. We applied the default HPF cut-off of 128 seconds. Contrast images were created for the safe and risky option regressors. We normalised the segmented anatomical images and the contrast images to a standard space (IXI549Space) coordinates, with a voxel size 1.5 ∗ 1.5 ∗ 1.5 *mm*. The contrast images were smoothed with a 6 *mm* FWHM isotropic Gaussian kernel.

During the second-level analyses, we performed a flexible factorial model with choice (safe, risky) and condition (baseline, experimental) as within-subject factors to examine state effects on risk-taking mechanisms. We compared the mean activity of risky choices to the mean activity of safe choices to analyse the main effect of choice. For the main effect of the condition, we compared the activity of both safe and risky choice in the baseline condition to the activity of both safe and risky choice in the experimental condition. Finaly, we analysed the interaction of choice and condition.

We compared the neural activity related to the different choices in the sated condition to those in the fasted conditions for study 1. For study 2, we compared the choice-related activity in the well-rested conditions to those in the sleep-deprived condition. We restricted our analyses to regions related to risk-taking and used therefore the different ROIs involved at the choice option onset from the mask of the meta-analysis of Cui and colleagues (2022) for small-volume corrections. These ROIs include bilateral ventral striatum, right dmPFC, and left middle occipital gyrus.

Next, we looked at the effect of ghrelin in each study. explained by the difference in ghrelin levels between conditions (baseline versus manipulation) on the interaction between condition and choice. We extracted six difference images from the first-level GLMs for each study: the difference between conditions for safe choices (baseline minus experimental and experimental minus baseline), the difference between conditions for risky choices, the interactions of choice and conditions. These difference images where then smoothed (6 *mm* FWHM), and correlated with the z-scored difference in ghrelin levels between conditions (baseline versus manipulation). We used the meta-analysis of Schulz and colleagues (2023; https://identifiers.org/neurovault.collection:13247) as mask for small-volume correction.

## Results

### Behavioural Results

In study 1, the median proportion of risky choices was descriptively higher in the sated condition (SAT; *Mdn_SAT_* = 0.45) compared to the fasted condition (FAS; *Mdn_FAS_* = 0.39; Figure 2.A). However, the data provided moderate evidence against a significant difference between conditions (*BF* = 0.31 ± 0.04%). In study 2, the data provided anecdotal evidence against condition differences in the mean proportion of risky choices (total sleep deprivation, TSD; night of normal sleep, NNS; *Mdn*_*TSD*_ = 0.44, *Mdn*_*NNS*_ = 0.38, *BF* = 0.52 ± 0.03% ; Figure 2.B). While reaction times were not the focus of our analysis, the data provided moderate (study 1; *BF* = 0.28 ± 0.04%) to strong (study 2; *BF* = 0.18 ± 0.05%) evidence against a significant difference between the baseline and the experimental conditions. Further analysis of reaction times can be found in the Supplementary material (Figure S1).

**Figure 2.**
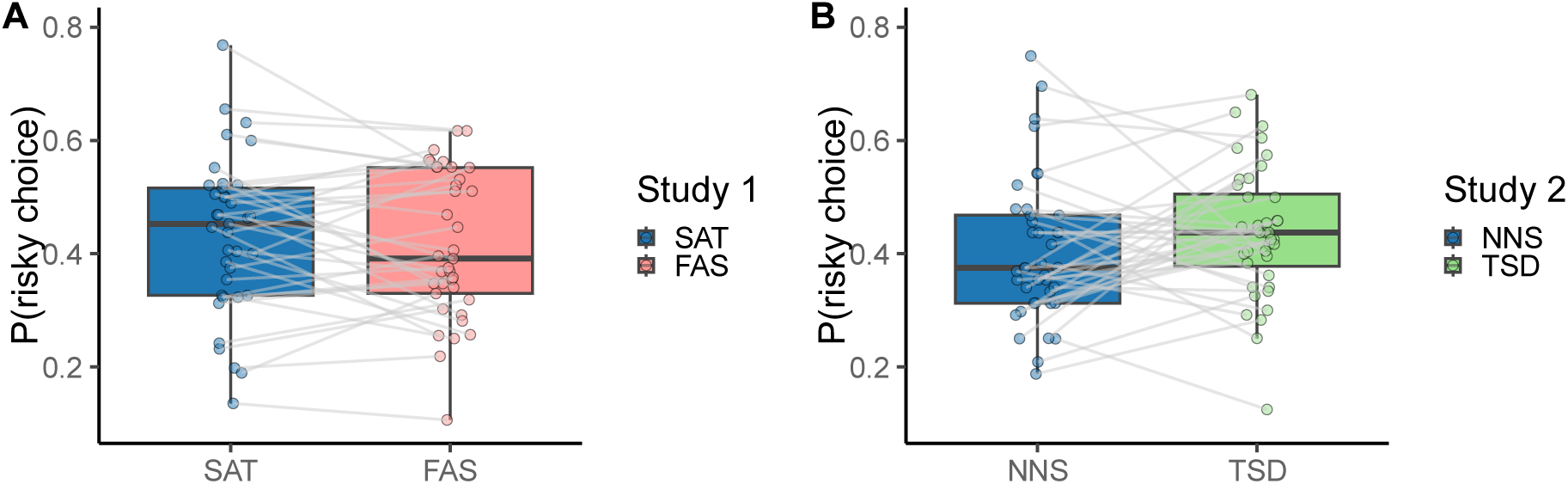
Median Proportion of Risky Choices. In study 1 (A), the median proportion of risky choices was higher in the sated condition (*Mdn_SAT_* = 0.45; blue), compared to the fasted condition (*Mdn_FAS_* = 0.39; red). However, the data provided moderate evidence against a significant difference between both conditions (*BF* = 0.31, ±0.04%). In study 2 (B), the median proportion of risky choices was smaller after a night of normal sleep (*Mdn_NNS_* = 0.38; blue), compared to one night of total sleep deprivation (*Mdn_TSD_* = 0.44; green). However, the data provided anecdotal evidence against a significant difference between both conditions (*BF* = 0.52 ± 0.03%).

Zooming in on the different reward probabilities, Bayesian ANOVA showed for study 1 that there was extreme evidence for an effect of reward probabilities on proportion of risky choices (*BF* = 1.4 ∗ 10^772^ ± 0.01%; see Figure S2.A in the Supplementary material) and anecdotal evidence against an effect of condition (*BF* = 0.07 ± 0.35%). There was evidence against an interaction effect of reward probability and condition (*BF* = 5.14 ∗ 10^−5^ ± 1.64%). In study 2, there was evidence for an effect of reward probabilities on proportion of risky choices (*BF* = 7.41 ∗ 10^394^ ± 0.01%) and anecdotal evidence against an effect of condition (*BF* = 0.35 ± 0.07%). There was very strong evidence against an interaction effect of reward probability and condition (*BF* = 0.01 ± 2.15%; see Figure S2.B in the Supplementary material). For the median reaction times, the data provided moderate evidence for an effect of reward probability in study 2 (*BF* = 4.28 ± 0.16) but not in study 1 (*BF* = 0.27 ± 0%). More details can be found in the Supplementary material.

### Endocrine Results

Ghrelin levels per study, subject and experimental condition are provided in Figure 3. The data demonstrated extreme evidence for an increase in ghrelin levels after fasting in study 1 (*Mdn_SAT_* = 269.52, *Mdn_FAS_* = 449.06, *BF* = 348838.8 ± 0%). In contrast, only anecdotal evidence for an increase in ghrelin levels following sleep deprivation was found in study 2 (*Mdn_NNS_* = 727.14, *Mdn_TSD_* = 769.3, *BF* = 2.17 ± 0%). Note that for study 2, ghrelin levels in the control condition were substantially greater than control condition ghrelin values in study 1. Rihm and colleagues (2019) used frequentist analyses, which yields a significant difference in ghrelin levels between normal sleep and a single night of total sleep deprivation. In contrast, Bayesian inference suggests that there is only inconclusive evidence for an increase in ghrelin levels.

**Figure 3.**
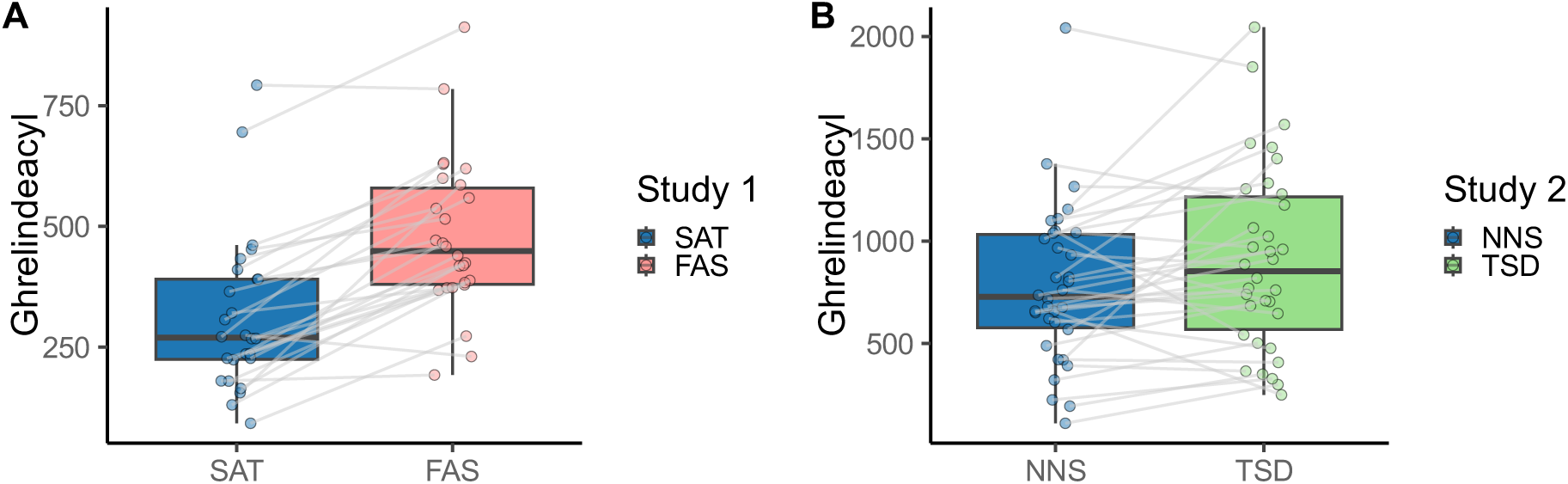
Ghrelin Levels. The ghrelin levels increased after fasting (A; *Mdn_SAT_* = 269.52, *Mdn_FAS_* = 449.06), it did not change after a single night of total sleep deprivation (B; *Mdn_NNS_* = 727.14, *Mdn_TSD_* = 769.3). However, note that ghrelin levels after normal sleep were already higher compared to the sated condition in study 1.

The effect of the manipulation on ghrelin levels disappeared when controlling for leptin, insulin, cortisol, and glucose in study 1. There was anecdotal evidence against a conditional effect on ghrelin (*BF* = 0.66 ± 0.69%). In study 2, the conditional effect remained absent (*BF* = 0.52 ± 3.64%). However, note that these peptides are not independent, violating the assumption of no multicollinearity (Figure S10 - Figure S15 in Supplementary Material). Nevertheless, these results suggest that there is no difference in ghrelin levels between the baseline and experimental conditions in both studies when controlling for the different peptides. The group differences of leptin, insulin, cortisol, and glucose are analysed in the Supplementary Material (Figure S9, Figure S16, Figure S20, and Figure S21 respectively).

To analyse the effect of ghrelin on the proportion of risky choices, we compared a Bayesian linear model with ghrelin and condition as predictors of the proportion of risky choices to a model with only condition as predictor. Both models included a random intercept for the participants. The data demonstrated anecdotal evidence in favour of the model with only condition as predictor of risky choices (*BF* = 0.44 ± 2.93%) in study 1. Similarly, there was anecdotal evidence against an effect of ghrelin on risky choices (*BF* = 0.62 ± 1.83%) in study 2.

### Model-Based Results

Results from the model comparison via WAIC showed that model variants including a valuation component (either PT or HPDM) were superior to the baseline models (see Figure S4 in the Supplementary material). Model fit was not improved by the inclusion of a perseveration term, neither FOP, nor HOP. Across studies and conditions, the PT model without perseveration provided the best account of the data. To verify that the best-fitting computational model provided a good account of observed behaviour, we performed posterior predictive checks (see Figure 4). Proportions of risky choices simulated from the PT model showed large overlap with the observed behaviour. This was the case across the full range of reward probabilities, and across studies and experimental conditions.

**Figure 4.**
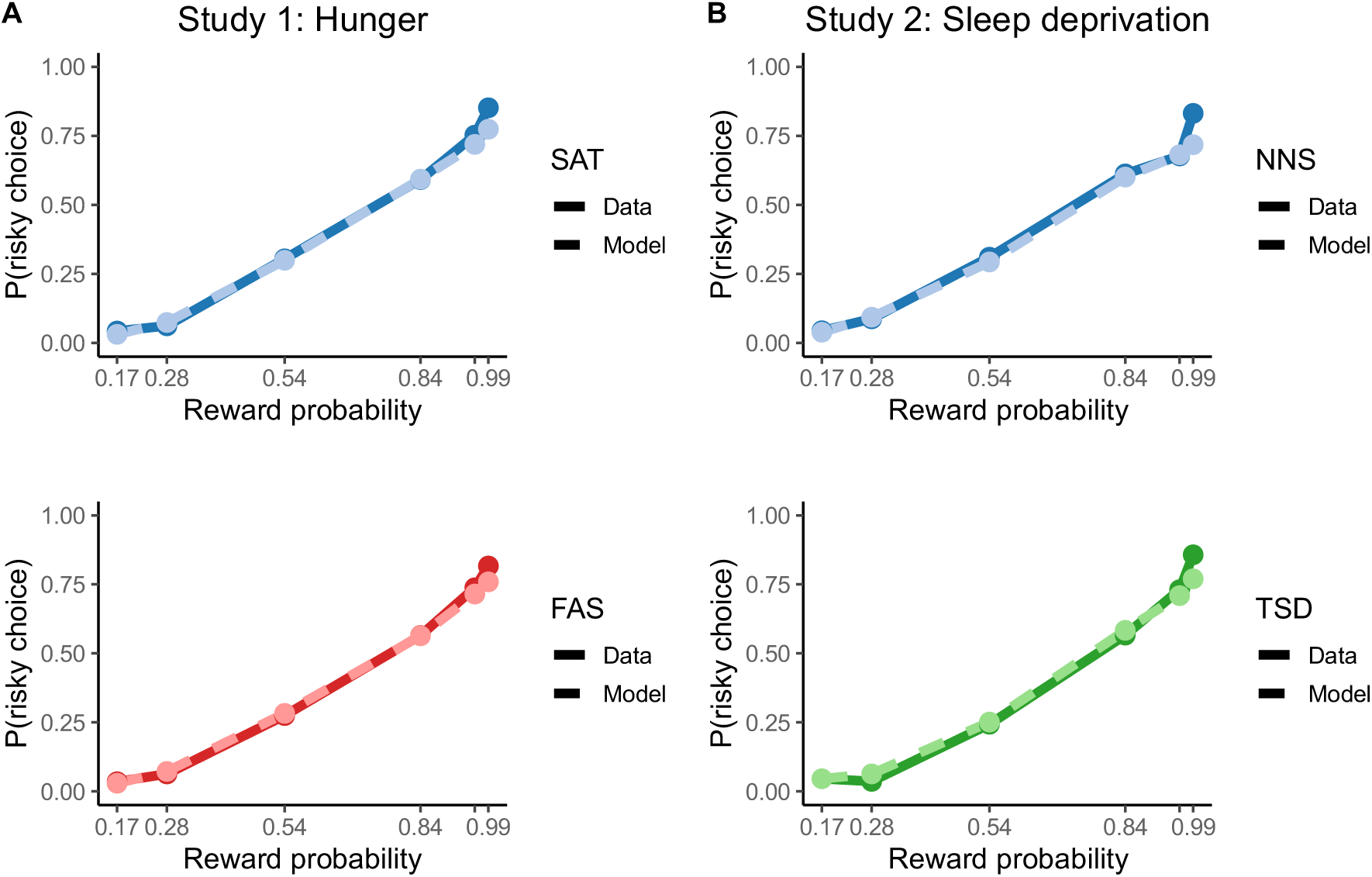
Choice Predictions and Behavioural Data. The probability of selecting the risky option per reward probability (0.17, 0.28, 0.54, 0.84, 0.96, 0.99) for participants (full line) and simulated by the prospect theory model (dashed line). The model predictions show a large overlap with the behavioural data. It is at the highest reward probability (*P* = 0.99) that the model diverges from the observation.

Next, we examined model parameters, focusing on the best-fitting PT model fitted to the full data set across experimental conditions (see Equation 8). The posterior distributions of the different parameters and their HDIs are displayed in Figure 5, and numerical values are provided in Table 2 and Table 3. In study 1, there were no relevant changes in parameter values between conditions at the group level (see Table 3). The 95% HDI contained zero for the difference distributions of each parameter. In study 2, there was a descriptive increase in attractiveness to risk (δ) and the policy parameter (β) after a single night of total sleep deprivation while sensitivity to probabilities (γ) decreased. However, the HDI for the difference distributions of each parameter in study 2 contained zero and we therefore conclude that there were no relevant changes in parameter values between conditions at the group level (see Table S2, Table S3, and Figure S5 in the Supplementary material for a replication with the subset of participants used for the fMRI analyses).

**Figure 5.**
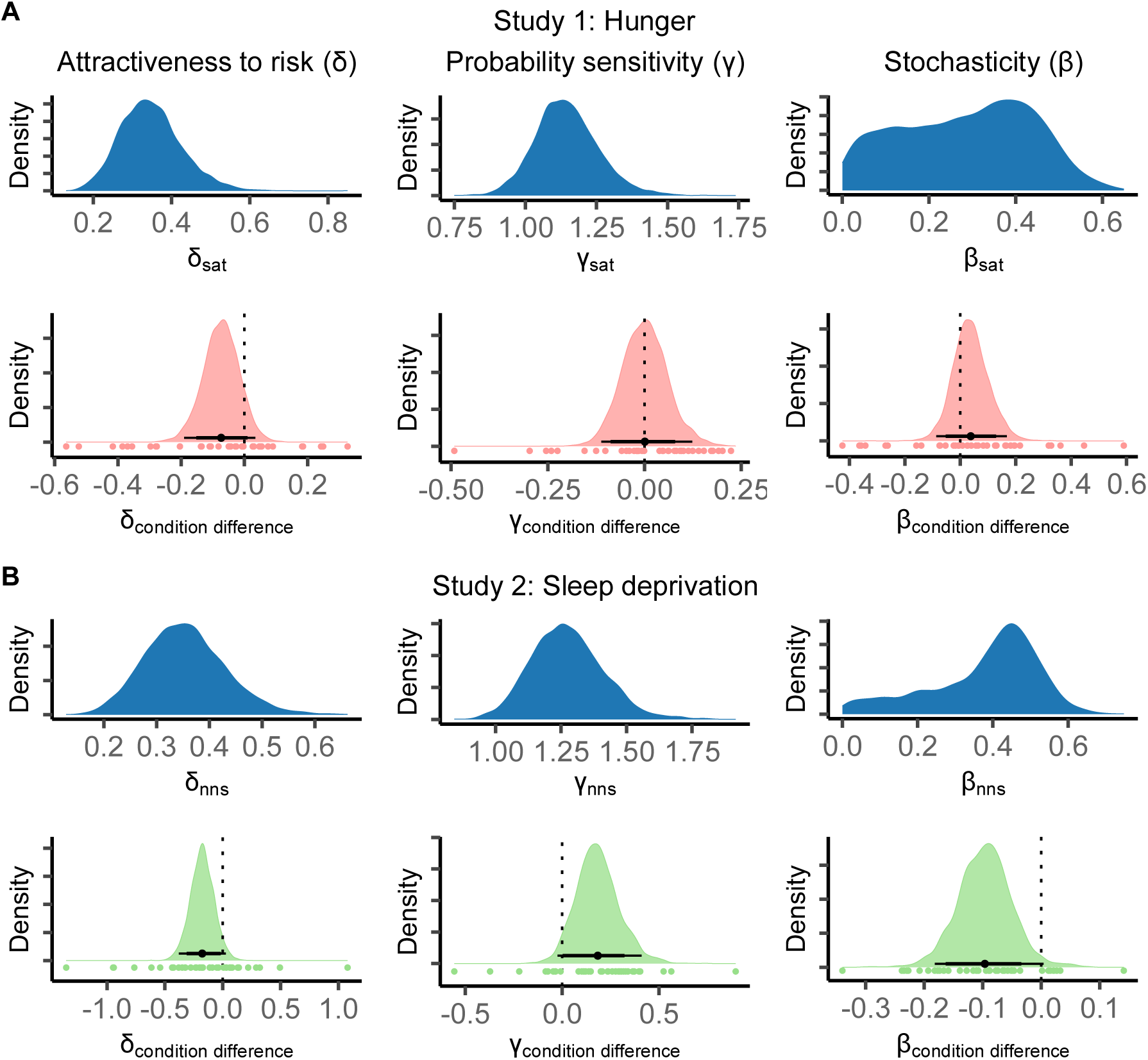
Posterior Distribution of Group-Level Parameters of the Prospect Theory Model. The posterior distributions of the group-level parameters for study 1 (hunger; A) and study 2 (sleep deprivation; B): attractiveness to risk δ, sensitivity to probabilities γ, policy parameter β. Parameter values of the sated (A) and rested (B) condition are shown in blue, the difference between conditions is shown in red for study 1 and in green for study 2. The horizontal lines represent the highest distribution interval (HDI) of 95% (thin line) and 85% (thick line), with the mean of the distribution as dot. The coloured dots underneath the distribution represent the mean values of individual subjects. See Table 2 for mean and SD values. All parameters, in both studies, did not differ significantly between conditions as shown by the HDI containing zero in each difference distribution.

**Table 2.** Prospect Theory Parameter Values. The mean (±SD) of the blue distributions shown in Figure 5: the estimated parameters for the PT model for the baseline condition (i.e., sated or night of normal sleep) and the experimental condition (i.e., fasted or total sleep deprivation) at the group level. Attractiveness to risk δ, sensitivity to probabilities γ, policy parameter β.

|  |  | Study 1 | Study 2 |
| --- | --- | --- | --- |
| $\delta$ | Baseline condition | 0.35 (±0.08) | 0.35 (±0.08) |
| | Experimental condition | 0.31 (±0.08) | 0.26 (±0.07) |
| $\gamma$ | Baseline condition | 1.15 (±0.11) | 1.27 (±0.14) |
| | Experimental condition | 1.15 (±0.12) | 1.49 (±0.18) |
| $\beta$ | Baseline condition | 0.28 (±0.15) | 0.38 (±0.15) |
| | Experimental condition | 0.32 (±0.16) | 0.28 (±0.15) |

**Table 3.** Condition Differences in Prospect Theory Parameters. The difference between the baseline and experimental condition for each study (the coloured distributions in Figure 5).

| | Mean (±SD) | 95% HDI | $dBF_{10}$ |
| --- | --- | --- | --- |
| Study 1 |  |  |  |
| Attractiveness to risk ($\delta$) | -0.07 (±0.06) | -0.19, 0.03 | 0.11 |
| Sensitivity to probabilities ($\gamma$) | 0 (±0.06) | -0.11, 0.12 | 1 |
| Policy parameter ($\beta$) | 0.04 (±0.06) | -0.1, 0.16 | 2.57 |
| Study 2 |  |  |  |
| Attractiveness to risk ($\delta$) | -0.18 (±0.1) | -0.38, 0.03 | 0.05 |
| Sensitivity to probabilities ($\gamma$) | 0.18 (±0.11) | -0.02, 0.4 | 23.26 |
| Policy parameter ($\beta$) | -0.09 (±0.05) | -0.18, 0 | 0.02 |

To investigate the effect of ghrelin on the parameters, we added ghrelin coefficients to the PT model as described in Equation 10. Note that the sample size is smaller since we only used the data of those participants with endocrine and neuroimaging data (see Table S4, Table S5, and Figure

S6 in the Supplementary material for the results of the models’ main parameters). The densities of the different ghrelin coefficients and their HDIs are displayed in Figure 6, their values in Table 4. In both study 1 and 2, there was no credible evidence for effects of ghrelin on any of the estimated parameters of the PT model.

**Figure 6.**
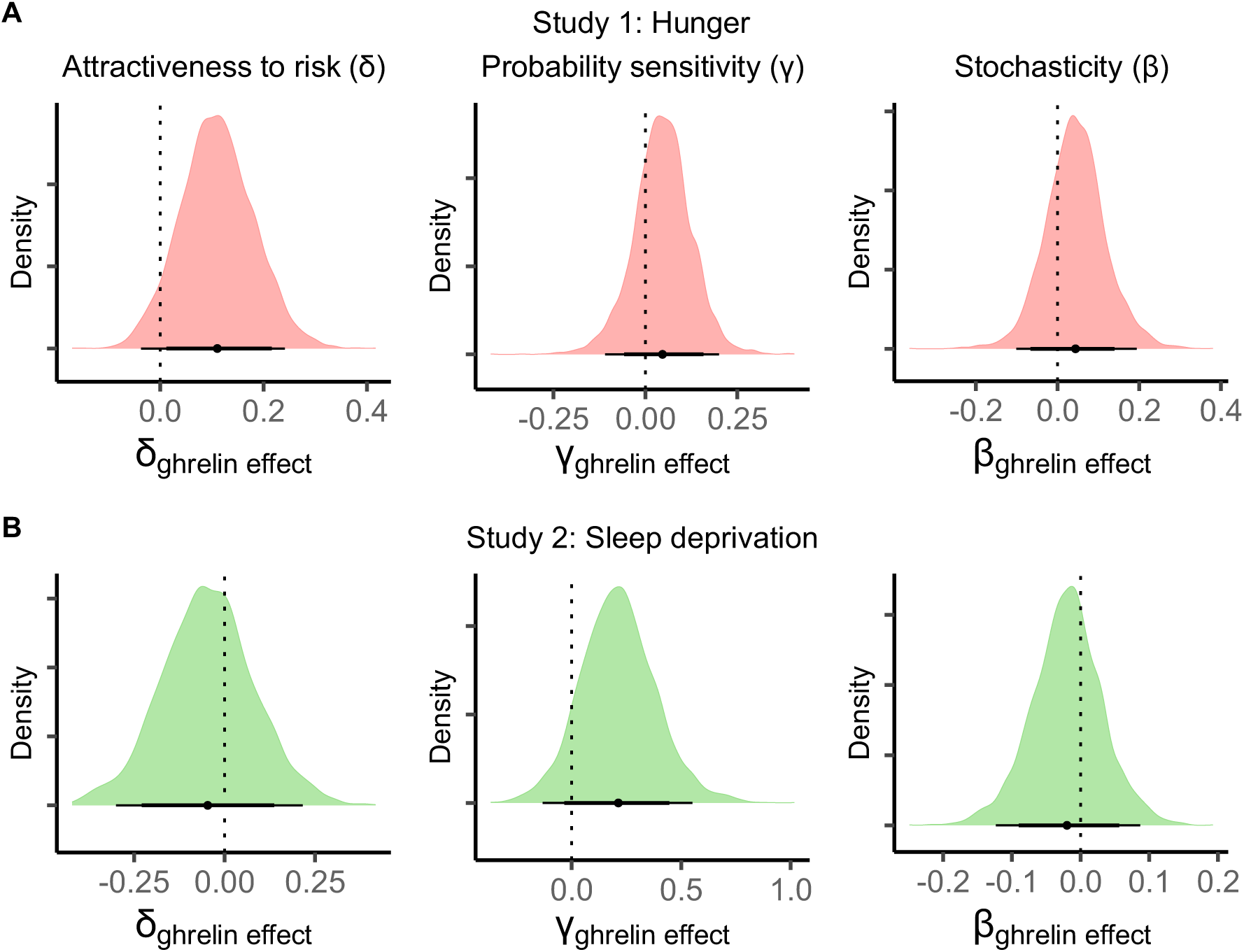
Posterior Distribution of Ghrelin Coefficients for the Parameters of the Prospect Theory Model. The posterior distributions of the ghrelin coefficients for the group-level parameters for study 1 (hunger; A) and study 2 (sleep deprivation; B): attractiveness to risk δ, sensitivity to probabilities γ, policy parameter β. Zero falls within the highest density intervals (HDI; thick bars: 0.85, thin bars: 0.95) for all parameter in both studies.

**Table 4.** Ghrelin Coefficients of the Prospect Theory Parameters. The effect of ghrelin on the estimated parameter values of the PT model presented in Figure 6.

| | Mean (±SD) | 95% HDI | $dBF_{10}$ |
| --- | --- | --- | --- |
| Study 1 |  |  |  |
| Attractiveness to risk ($\delta$) | 0.11 ($\pm 0.07$) | −0.04, 0.24 | 15.61 |
| Sensitivity to probabilities ($\gamma$) | 0.05 ($\pm 0.08$) | −0.11, 0.2 | 2.86 |
| Policy parameter ($\beta$) | 0.04 ($\pm 0.07$) | −0.1, 0.19 | 2.79 |
| Study 2 |  |  |  |
| Attractiveness to risk ($\delta$) | −0.05 ($\pm 0.13$) | −0.3, 0.22 | 0.55 |
| Sensitivity to probabilities ($\gamma$) | 0.21 ($\pm 0.17$) | −0.13, 0.55 | 8.94 |
| Policy parameter ($\beta$) | −0.02 ($\pm 0.05$) | −0.12, 0.09 | 0.52 |

## Neuroimaging Results

### Subjective Value Effects

The subjective value was calculated with the mean value of the estimated parameter distributions of each participant. To analyse the activity related to subjective value we used the ROIs from the mask of the Rangel laboratory, FWE-SVC (see Methods section). Activity of subjective value is in line with previous studies (Figure 7, see Table S6 in the Supplementary material for an overview of both studies and the MNI coordinates of the regions with statistically significant activity). Study 1 showed a main effect of subjective value in the right PCC, right ventral striatum (NAc), bilateral ACC, and left nucleus caudatus. In study 2, a main effect of subjective value was found in the left nucleus caudatus and ACC, and right PCC. Across both studies, none of the BOLD differences between conditions survived the stricter threshold of FWE-based small volume correction. Additionally, we did not find any region where conditional differences of ghrelin correlated significantly with the neural signatures of subjective value. This, again, was true across both studies.

**Figure 7.**
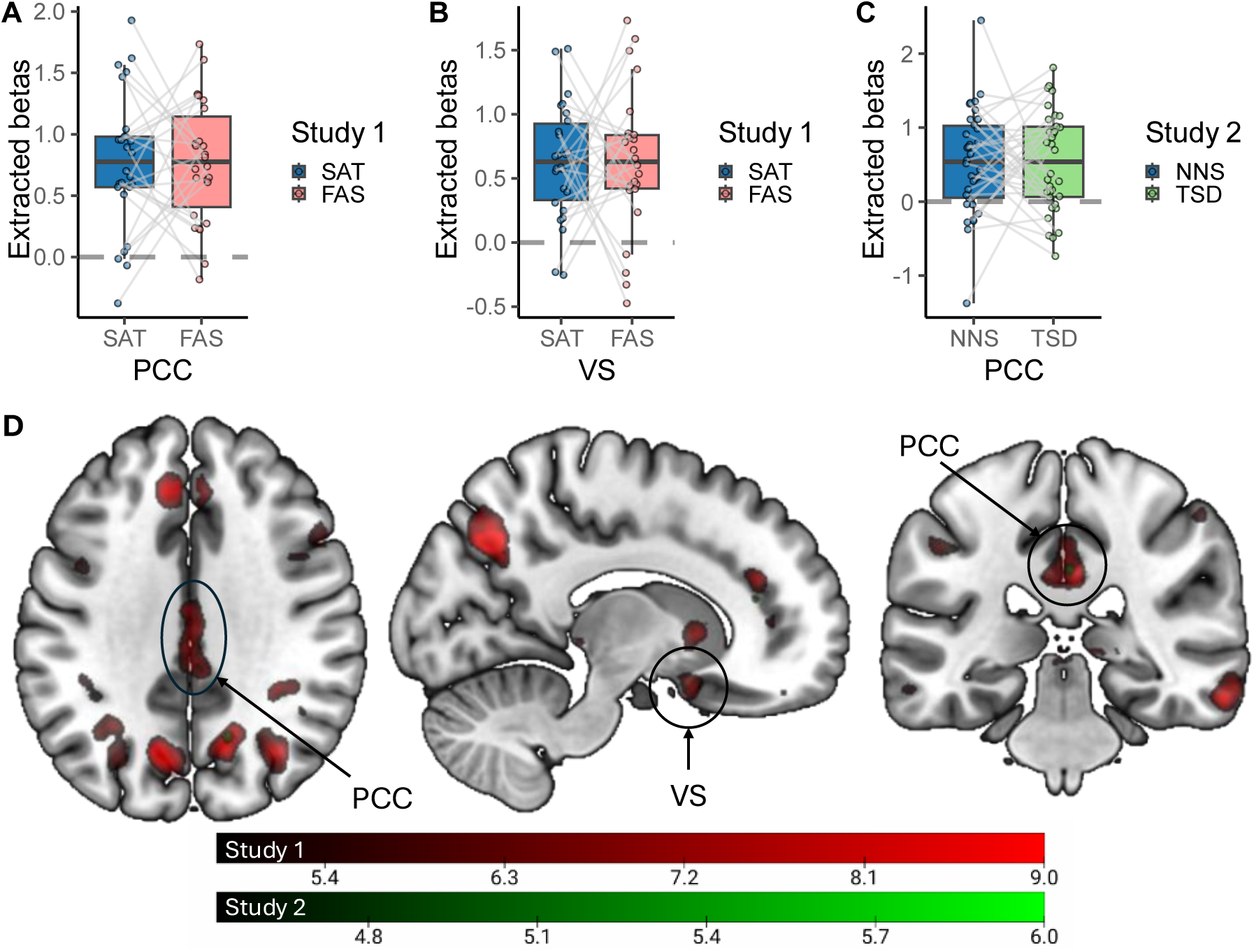
Neural Activity Related to Subjective Value. A-C) The extracted β-coefficients of subjective value in the right PCC (A; *X* = 2, *Y* = −32, *Z* = 38) and VS (B; 12, 11, −14) for study 1, and the right PCC (C; 2, −32, 34) for study 2. The β-coefficients support the finding of a main effect of subjective value but no effect of condition. D) Clusters with significant activity for subjective value for study 1 (red) and study 2 (green). Slices, made on *X* = 13, *Y* = −32, *Z* = 34, support the overlap between studies. However, the activity in study 2 is smaller than in study 1. Display threshold: *p* < .001 uncorrected.

### Choice Effects

We used the ROIs from the mask of the meta-analysis of Cui and colleagues (2022), FWE-SVC, for the analysis of neural activity related choices (see Methods section). Study 1 showed increased activity for risky choices compared to safe choices in the right dlPFC, right inferior parietal gyrus, right inferior temporal gyrus, left anterior insula, and bilateral PCC. Study 2 showed increased activity in the left nucleus caudatus and right dlPFC for risky choices (full results are reported in Table S7 and Figure S7 in the Supplementary material). There was no increased activity for safe choices that survived the threshold in either study.

No region showed significant activity related to the interaction of choice (risky versus safe) and condition in our a prior defined ROIs, neither in study 1 nor study 2. Additionally, we did not find any region where conditional differences of ghrelin correlated significantly with the neural activity of choice or the activity of the interaction of choice and condition. This, again, was true across both studies.

## Discussion

In two separate studies, we manipulated ghrelin levels via short-term fasting (study 1) or a single night of total sleep deprivation (study 2), to test the prediction that increases in ghrelin levels would be linked to increased risk-taking (Dickinson et al., 2022; Levy et al., 2013; Pietrzak et al., 2023; Shabat-Simon et al., 2018; Venkatraman et al., 2007, 2011). Bayesian analyses revealed that the fasting manipulation successfully elevated ghrelin levels, whereas the evidence for an effect of sleep deprivation on ghrelin levels was inconclusive (however, see Rihm et al., 2019 for a significant effect using frequentist analysis on the same data). In contrast with some previous findings (Brunet et al., 2020; Shabat-Simon et al., 2018; van Swieten et al., 2023; Venkatraman et al., 2011), neither manipulation affected model-agnostic or computational modelling based measures of risky decision-making. There was no credible evidence for an effect of ghrelin levels on risky choice behaviour or model parameters.

Neither experimental manipulation reliably affected the proportion of risky choices, which is in line with some previously reported null effects (Mao et al., 2023; van Swieten et al., 2023). In contrast, several studies reported an increase in risk-taking after fasting (Levy et al., 2013; Shabat-Simon et al., 2018; Symmonds et al., 2010) or sleep deprivation (Brunet et al., 2020; Dickinson et al., 2022; Mckenna et al., 2007; Salfi et al., 2020; Womack et al., 2013).

Fasting and sleep deprivation are distinct processes that may affect decision-making and risk-taking via different routes (Harrison & Horne, 2000; Heisler & Lam, 2017; Krause et al., 2017; Liu & Kanoski, 2018). Hunger interacts, via the hippocampus (Davidson et al., 2007; Kanoski & Grill, 2017) and hypothalamus (Siemian et al., 2021; Wallner-Liebmann et al., 2010), with the VTA and NAc of the dopaminergic reward network (Cassidy & Tong, 2017; Kahn & Shohamy, 2013; Nieh et al., 2016). Moreover, hunger may increase the connectivity between the hypothalamus and the dlPFC (Kullmann et al., 2023). Sleep deprivation may decrease thalamic and vmPFC activity (Menz et al., 2012; Thomas et al., 2000), and it may decrease the ability to discriminate between different rewards by increasing general reward processing in the mesolimbic system (Krause et al., 2017).

Importantly, however, both fasting and sleep deprivation elicit effects within the dopaminergic reward network. One possible element that may explain some of the overlapping effects is ghrelin (Kulkarni et al., 2024; Müller et al., 2015; Schulz et al., 2023; Stievenard et al., 2017). VTA and NAc, core regions of the DA system, are modulated by ghrelin (Abizaid et al., 2006; Al Massadi et al., 2019; Schulz et al., 2023; Stievenard et al., 2017). Since ghrelin increases both following fasting and sleep deprivation (AlDabal, 2011; Cummings et al., 2001; Müller et al., 2015; Schmid et al., 2008) it has been hypothesized as a potential mechanism underlying effects on risky-decision making (Müller et al., 2015; Schulz et al., 2023; Sztainert et al., 2018). However, we did not find credible evidence for associations between ghrelin levels and risk-taking.

Another peptide that is linked to both hunger and sleep deprivation is leptin (Austin & Marks, 2009; Klok et al., 2007; Spiegel, Leproult, et al., 2004; van Egmond et al., 2023). Although leptin signals energy levels (Ahima & Flier, 2000; Friedman, 2019), it does not fluctuate significantly between meals (Austin & Marks, 2009; Friedman, 2019; Korbonits et al., 1997). In addition, while leptin levels decrease after restricted sleep during multiple days (Spiegel, Leproult, et al., 2004; Spiegel, Tasali, et al., 2004), it does not appear to drop after one night of total sleep deprivation in men (Schmid et al., 2008; van Egmond et al., 2023). Comparison of the median leptin levels in both studies confirmed that the present manipulations did not affect leptin levels (see Figure S9 in the Supplementary material), supporting the exclusion of leptin from the analyses. Nevertheless, its involvement in several state-dependent processes and its interaction with the VTA (Fulton et al., 2006; Geisler & Hayes, 2023; Opland et al., 2010), make leptin an interesting peptide for future studies on the state-mediated effects on risky decision-making.

Ghrelin and leptin are not the only peptides that may be affected by the experimental manipulations applied in this project. Fasting and sleep deprivation may each influence a distinct set of peptides. Hunger is, for example, paired with lower glucagon-like-peptide-1 (GLP-1) levels, a peptide that signals satiation (Elliott et al., 1993; Krieger, 2020). Both ghrelin and GLP-1 interact with the reward network, however, in opposite directions (Decarie-Spain & Kanoski, 2021; Schulz et al., 2023). Sleep deprivation affects several peptides and hormones (Kim et al., 2015; van Cauter et al., 2007), and modulates adenosine levels (Basheer et al., 2004; Elmenhorst et al., 2007), which may explain decreased cognitive performance following sleep loss. However, little is known about adenosine effects on higher cognitive functions such as risky decision-making. More research is needed to better understand the role of the different peptides and hormones that are affected by hunger and sleep deprivation.

While we expected increased risk-taking behaviour in the experimental conditions, we are not the first to report a lack of change in risk-taking following fasting (van Swieten et al., 2023) or sleep deprivation (Greer et al., 2016; Mao et al., 2023; Menz et al., 2012). It is possible that the results from earlier studies may reflect false positive effects due to small sample sizes (e.g., Brunet et al., 2020; de Ridder et al., 2014). However, mixed results were found with both smaller (Acheson et al., 2007; Brunet et al., 2020) and larger sample sizes (Mao et al., 2023; Sztainert et al., 2018). Another possible explanation for the inconclusive results in the literature may be related to the distinction between risk and uncertainty. In experimental designs such as the one applied here, participants are fully informed about reward probabilities (e.g., the probability is presented on the screen as in Figure 1) while the reward probability is unclear in other paradigms (Fox & Poldrack, 2009; Vives et al., 2023). Similarly to our results, paradigms with fully transparent probability information showed no effects of hunger (van Swieten et al., 2023) or ghrelin (Pietrzak et al., 2023), while paradigms with unclear probability information (where participants did not know the reward probabilities) tended to observe increased risk-taking when participants were fasted (Sztainert et al., 2018; van Swieten et al., 2023) or ghrelin was administered (Pietrzak et al., 2024). Another way of interpreting the difference in paradigm is the presence of learning. In tasks without transparent probability information, probabilities must be inferred from feedback. It is possible that ghrelin affects risky decision-making indirectly by modulating this learning process (van Swieten et al., 2021). Future research could explore this in more detail.

To investigate the potential effects of the state-dependent manipulations on the underlying processes, we followed a computational modelling approach. Our model comparison was consistent across studies and experimental conditions, and revealed that the PT model without perseveration provided the best account of the data. WAIC scores (Watanabe, 2010) penalise models with more parameters, thus decreasing the likelihood of overfitting (Vandekerckhove et al., 2015). Model comparison was also supported by posterior predictive checks, which confirmed that the best-fitting model accurately reproduced the observed data across reward probabilities and experimental conditions.

In contrast to previous work (Palminteri, 2023; Wiehler et al., 2021), accounting for perseveration did not further improve model fit. One likely explanation is the absence of exploration and exploitation in the current paradigm. Choice repetition may play a greater role in situations where participants need to find an optimal balance between exploration and exploitation (Abir et al., 2024; Lerner, 2020; Miller et al., 2019), e.g., during reinforcement learning in volatile environments perseveration may help to stabilize choice patterns (Ferguson et al., 2023; Wiehler et al., 2021). However, in the present risky choice paradigm, exploration is not required, as all choice-relevant information is provided.

Examination of parameter estimates derived from the best-fitting model allowed us to analyse latent processes involved in risky decision-making. Although there was no difference in risky choice behaviour between the baseline and experimental condition, it is possible that similar behavioural patterns can result from different combinations of parameter values. However, in line with the model-agnostic analyses, these analyses revealed little credible evidence for state-dependent effects. These null-results were relatively robust, and observed when analysing all participants, only the subset of participants completing the fMRI sessions, or when incorporating ghrelin as a covariate directly in the model. The only credible condition effect reflected an increase in choice stochasticity (reduction in the policy parameter beta) following one night of total sleep deprivation. This is in line with previous work (Glass et al., 2011; Menz et al., 2012), and confirms that sleep deprivation can lead to increases in choice stochasticity. Our results would be in line with the idea that sleep deprivation could have effects that are not elicited by fasting, which may drive increases in choice stochasticity. Sleep deprivation is strongly associated with stress and metabolism (AlDabal, 2011; Hirotsu et al., 2015; Kim et al., 2015; Kulkarni et al., 2024), and alters reward processing (Mullin et al., 2013; Rihm et al., 2019; Womack et al., 2013). Such effects might underlie the observed effects on choice stochasticity. Additionally, animal models suggest that adenosine levels increase in the basal forebrain during sleep deprivation (Basheer et al., 2004, 2007; Blanco-Centurion et al., 2006). The basal forebrain is involved in the control of attention (Maness et al., 2022) and increased levels of adenosine lower performance on attention tasks (Christie et al., 2008; Urry & Landolt, 2014). Decreased attention might provide an alternative explanation for the increased choice stochasticity (e.g., Hunt et al., 2018) after one night of total sleep deprivation. Note that, according to the Bayesian analysis, the evidence for increased ghrelin levels between the baseline and experimental conditions was inconclusive in study 2. This would be in line with the idea that other sleep deprivation related effects (and not ghrelin levels per se) may underlie the observed increases in choice stochasticity.

The small effect of one night of total sleep deprivation on choice stochasticity, but not on risky decision-making, could also potentially be linked to the decreased activity in the thalamus and PFC. The results of studies where participants underwent one night of total sleep deprivation are mixed. Some are in line with our results and did not observe an effect on risk-taking (Dickinson et al., 2022; Mao et al., 2023; Maric et al., 2017; Menz et al., 2012), while others found that participants showed increased risk-taking when sleep deprived (Mckenna et al., 2007; Venkatraman et al., 2011). Some authors state that sleep deprivation does not affect more complex cognitive processes (Goel et al., 2009; Harrison & Horne, 2000) and one could argue that risky decision-making would be one of them. However, studies with partial sleep deprivation over a longer period did find a positive effect on increased risky decision-making (Brunet et al., 2020; Maric et al., 2017; Salfi et al., 2020). Activity in the thalamus and PFC showed a continued decrease with increased periods of sleep deprivation (Thomas et al., 2000, 2003). It is possible that neural activity did not decrease sufficiently for it to affect risk-taking. Future research could explore the effects of longer periods sleep deprivation on risky decision-making.

There was also no credible evidence for an effect of condition differences in ghrelin levels on model parameters in either study. This adds to the ongoing discussion on metabolic influences on risk-taking, by showing that changes in ghrelin levels are unlikely to drive changes in risk-taking following two different metabolic manipulations.

The present study replicated several core neuroimaging effects. First, across both studies, using an a priori defined ROI approach, we replicated subjective value effects in the ACC, PCC, caudate, and ventral striatum (Bartra et al., 2013; Levy et al., 2010; Nachev et al., 2015; Pearson et al., 2011). Second, using a meta-analysis based mask (Cui et al., 2022), we replicated risky versus safe option choice effects in the anterior insula in study 1 and the caudate in study 2, in addition to the dlPFC across both studies. However, in line with the behavioural and modelling null-results, there were no effects of fasting or sleep deprivation on the neural encoding of subjective value or risky versus safe choices. Nor was there evidence for a modulatory effect of ghrelin on the neural encoding of subjective value or choices.

Several limitations of the present approach deserve mention. First, the samples consisted of only male participants, limiting generalizability. Not only do the different sexes show differences in risk-taking (Byrnes et al., 1999; Cornwall et al., 2018; van den Bos et al., 2013), but sex differences have also been observed in the effects of sleep deprivation (Acheson et al., 2007; Ferrara et al., 2015; Lim et al., 2022), and ghrelin effects on risk-taking (Pietrzak et al., 2023). Future research is needed to continue the investigation of sex differences in ghrelin effects. Second, sample sizes were relatively small (Grady et al., 2021; Poldrack et al., 2017), and future studies would benefit from larger sample sizes. Nevertheless, we replicated several fMRI effects in pre-registered ROIs, e.g., with respect to subjective value (Menz et al., 2012; Peters & Büchel, 2009; Seaman et al., 2018; Wang et al., 2023) and risky choice (Cui et al., 2022; Rao et al., 2008), showing sufficient power for replications. Third, modelling conclusions are restricted to the models included in our model space. We focused on the basic version of two models that have been successfully applied in previous studies, and posterior predictive checks confirmed that the models accurately reproduced observed choice patterns. However, it is possible that models may not have accurately captured the processes underlying behaviour in some participants (Levy et al., 2013). Therefore, it is possible that ghrelin did have an effect on latent choice variables which were not accounted for in the models we used in our analysis. Future research may include, for example, versions of the cumulative prospect theory (Fennema & Wakker, 1997; Harrison & Swarthout, 2023). Additionally, since we did not focus on reaction times, the inclusion of drift diffusion models could provide additional information about the underlying choice processes (Lee et al., 2023; Peters et al., 2020) and lead to potential interesting further insights.

In summary, we report on two independent fMRI studies which tested the degree to which state-dependent changes (in particular ghrelin levels) impacted risky decision-making. Participants performed a probability discounting task, either after a short fasting period or one night of total sleep deprivation. There were no effects of the experimental manipulations or increased ghrelin levels on risky choice behaviour or model-based measures. Finally, there was no neural activity related to the experimental manipulation or the increased ghrelin levels. This suggests that the state-dependent influences, in particular from ghrelin, on risky decision-making are weaker than previously thought.

## Acknowledgements

S.G. analysed the data. S.G. and J.P. wrote the paper. A.B. and H.S. provided analytical tools. J.P. and H.S. designed the study. All authors provided revisions. We would like to thank Julia Rihm for the data collection. Correspondence concerning this article should be addressed to Steven Geysen (sgeysen[at]uni-koeln[dot]de).

## Author note

We have no known conflict of interest to disclose.

## Data and code availability

The data and the code for this project are publicly available on OSF (https://osf.io/gcjeq/).

