## Supplementary Material for "Ghrelin and risky decision-making: No credible evidence for homeostatic state modulation of neural or behavioural effects"

#### Reaction times

The median reaction times (RTs) did not differ between conditions according to a paired Bayesian t-test ($BF = 0.18 \pm 0.05\%$; Figure S1). To analyse the effect of ghrelin on RTs, we compared a Bayesian linear model with ghrelin and condition as predictors of RTs to a model with only condition as predictor. Both models included a random intercept for the participants. The data demonstrated anecdotal evidence in favour of the model with only condition as predictor of RT ($BF = 0.92 \pm 3.77\%$) in study 1. In study 2, there was strong evidence in favour of an effect of ghrelin on RT ($BF = 22.09 \pm 10.91\%$).

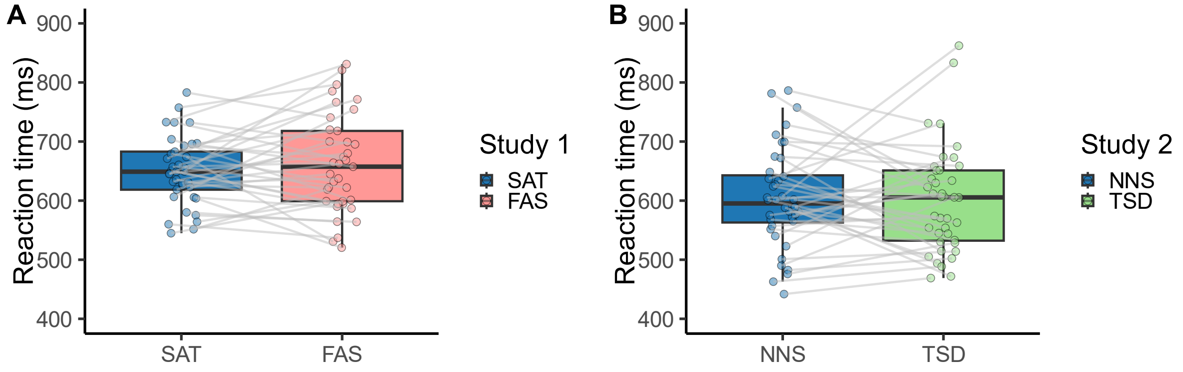


**Figure S1.** Median Reaction Times. Participants in study 1 (A) had faster median reaction times in the sated condition ($Mdn_{SAT} = 649\ ms$; blue), compared to the fasted condition ($Mdn_{FAS} = 657.5\ ms$; red). However, the data provided moderate evidence against a significant difference between both conditions ($BF = 0.28 \pm 0.04\%$). Median reaction times in study 2 (B) were faster after a night of normal sleep ($Mdn_{NNS} = 595.25\ ms$; blue), compared to one night of total sleep deprivation ($Mdn_{TSD} = 605.5\ ms$; green). The data provided strong evidence against a significant difference between both conditions ($BF = 0.18 \pm 0.05\%$).

**Effect of Reward Probability**

We analysed the proportion of risky choices and median reaction times (RTs) more closely by investigating the effect of reward probability and its interaction with condition. See Figure S2 and Figure S3 respectively.

We used a Bayesian ANOVA to analyse the effects of reward probability and condition. The results for the proportion of risky choices are presented in the main body of the paper as this was our focus. The results of the median RTs are presented in more detail here. In study 1, there was moderate evidence against an effect of reward probability ($BF = 0.29 \pm 0\%$) and condition ($BF = 0.14 \pm 0.18\%$) on median RT. There was extreme evidence against an interaction effect of reward probability and condition ($BF = 7.3 * 10^{-5} \pm 5.24\%$). In study 2, the data provided moderate evidence for an effect of reward probability on median RT ($BF = 4.28 \pm 0.16\%$) and anecdotal evidence against an effect of condition on median RT ($BF = 0.08 \pm 0.3\%$). There was extreme evidence against an interaction effect of reward probability and condition ($BF = 0 \pm 3.23\%$).

The uncorrected results of Bayesian paired t-tests per reward probability are presented in Table S1.

**Table S1.** Bayes Factor per Reward Probability. Results from Bayesian paired t-test ($BF \pm$ proportional error estimate) comparing the difference in the proportion of risky choices from the baseline condition (i.e., sated or night of normal sleep) and the experimental condition (i.e., fasted or total sleep deprivation) to zero, per reward probability. The data did not contain any evidence of such differences. The same is true for reaction times. Uncorrected for multiple comparisons.

| Reward probability | 0.17 | 0.28 | 0.54 | 0.84 | 0.96 | 0.99 |
|---|---|---|---|---|---|---|
| Study 1 | | | | | | |
| Proportion of risky choices | | | | | | |
| | 0.08 $\pm$0.27% | 0.07 $\pm$0.31% | 0.12 $\pm$0.17% | 0.08 $\pm$0.25% | 0.08 $\pm$0.27% | 0.21 $\pm$0.1% |
| Reaction time | | | | | | |
| | 0.08 $\pm$0.27% | 0.07 $\pm$0.31% | 0.1 $\pm$0.2% | 0.15 $\pm$0.14% | 0.24 $\pm$0.09% | 0.07 $\pm$0.31% |
| Study 2 | | | | | | |
| Proportion of risky choices | | | | | | |
| | 0.09 $\pm$0.2% | 1.01 $\pm$0.2% | 0.6 $\pm$0.03% | 0.1 $\pm$0.18% | 0.21 $\pm$0.09% | 0.14 $\pm$0.13% |
| Reaction time | | | | | | |
| | 0.09 $\pm$0.2% | 0.11 $\pm$0.17% | 0.31 $\pm$0.06% | 0.09 $\pm$0.2% | 0.31 $\pm$0.06% | 0.1 $\pm$0.18% |

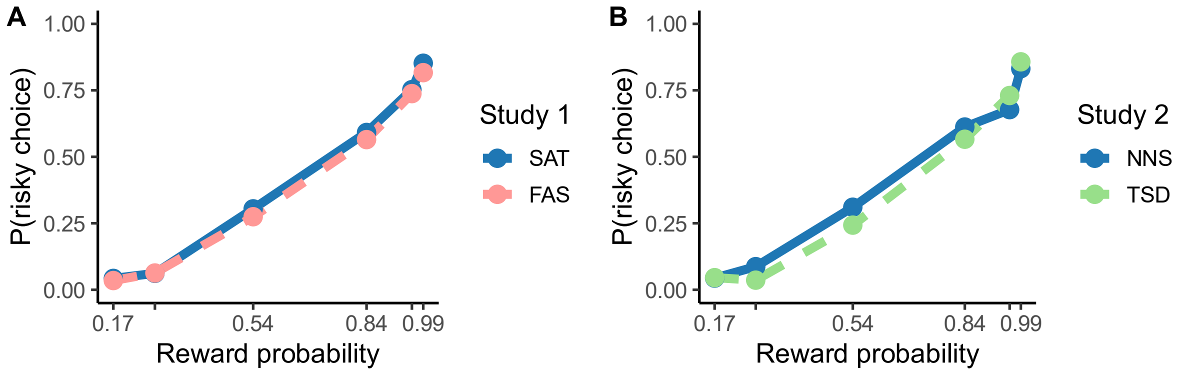


**Figure S2.** Proportion of Risky Choices per Reward Probability. The probability of selecting the risky option per reward probability $(0.17, 0.28, 0.54, 0.84, 0.96, 0.99)$ for each condition.

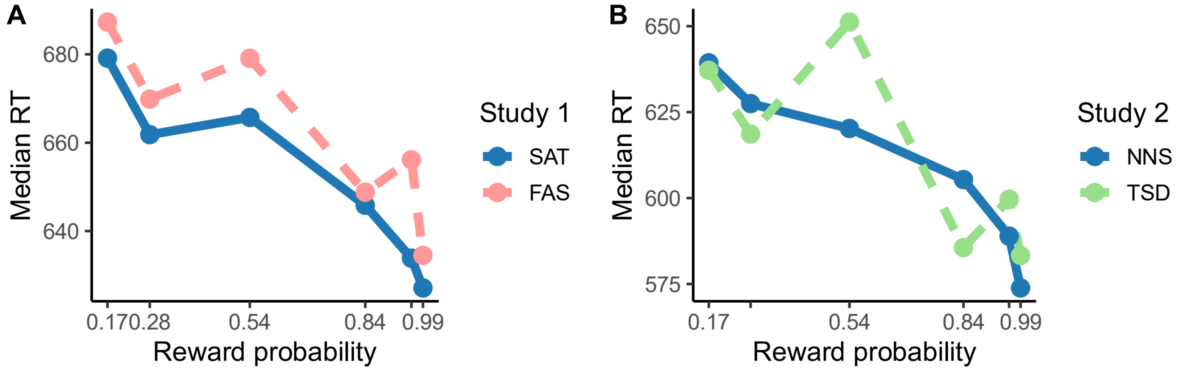


**Figure S3.** Median Reaction Times per Reward Probability. The median reaction times per reward probability $(0.17, 0.28, 0.54, 0.84, 0.96, 0.99)$ for each condition.

### Model-Based Results

#### *Model Comparison*

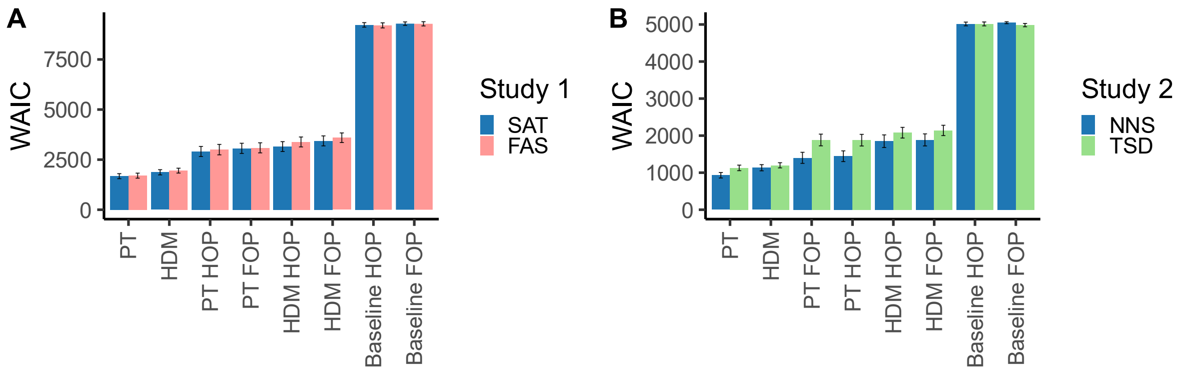


**Figure S4.** WAIC Scores of Model Space. The mean WAIC scores for each model per condition (error bars show the standard error). Both the prospect theory (PT) model and hyperbolic probability discounting model (HDM) without perseveration terms performed better than the other models. Since PT had the lowest WAIC score in both conditions, in both studies, we interpreted it as the best fitting model and continued our analyses with this model.

***Prospect Theory with fMRI Participants***

Here (Table S2, Table S3, and Figure S5) we report the parameter values of the PT model fitted to the subset of participants of which the neuroimaging data was analysed. The values differ from those in Table 3 and Table 4 due to the different sample sizes. However, the interpretation of the results does not change.

**Table S2.** Prospect Theory Parameter Values with fMRI Participants. Mean (±SD) of the blue distributions shown in Figure S5. Estimated parameters for the PT model for the baseline condition (i.e., sated or night of normal sleep) and the experimental condition (i.e., fasted or total sleep deprivation) at the group level. The model was fitted to the subset of participants of which the neuroimaging data was analysed. Attractiveness to risk $\delta$, sensitivity to probabilities $\gamma$, policy parameter $\beta$.

| | | Study 1 | Study 2 |
|---|---|---|---|
| $\delta$ | Baseline condition | $0.32\ (\pm 0.08)$ | $0.34\ (\pm 0.08)$ |
| | Experimental condition | $0.3\ (\pm 0.08)$ | $0.24\ (\pm 0.08)$ |
| $\gamma$ | Baseline condition | $1.2\ (\pm 0.15)$ | $1.32\ (\pm 0.17)$ |
| | Experimental condition | $1.25\ (\pm 0.16)$ | $1.54\ (\pm 0.22)$ |
| $\beta$ | Baseline condition | $0.27\ (\pm 0.17)$ | $0.35\ (\pm 0.14)$ |
| | Experimental condition | $0.24\ (\pm 0.18)$ | $0.24\ (\pm 0.15)$ |

**Table S3.** Conditional Differences of Prospect Theory with fMRI Participants. The difference between the baseline and experimental condition for each study (the coloured distributions in Figure S5).

| | Mean (±SD) | 95% HDI | $dBF_{10}$ |
|---|---|---|---|
| Study 1 | | | |
| Attractiveness to risk ($\delta$) | $-0.05$ ($\pm 0.06$) | $-0.18, 0.08$ | 0.27 |
| Sensitivity to probabilities ($\gamma$) | 0.04 ($\pm 0.06$) | $-0.08, 0.18$ | 3.23 |
| Policy parameter ($\beta$) | $-0.03$ ($\pm 0.07$) | $-0.17, 0.12$ | 0.45 |
| Study 2 | | | |
| Attractiveness to risk ($\delta$) | $-0.2$ ($\pm 0.12$) | $-0.41, 0.05$ | 0.05 |
| Sensitivity to probabilities ($\gamma$) | 0.19 ($\pm 0.14$) | $-0.08, 0.45$ | 11.4 |
| Policy parameter ($\beta$) | $-0.11$ ($\pm 0.05$) | $-0.2, 0$ | 0.02 |

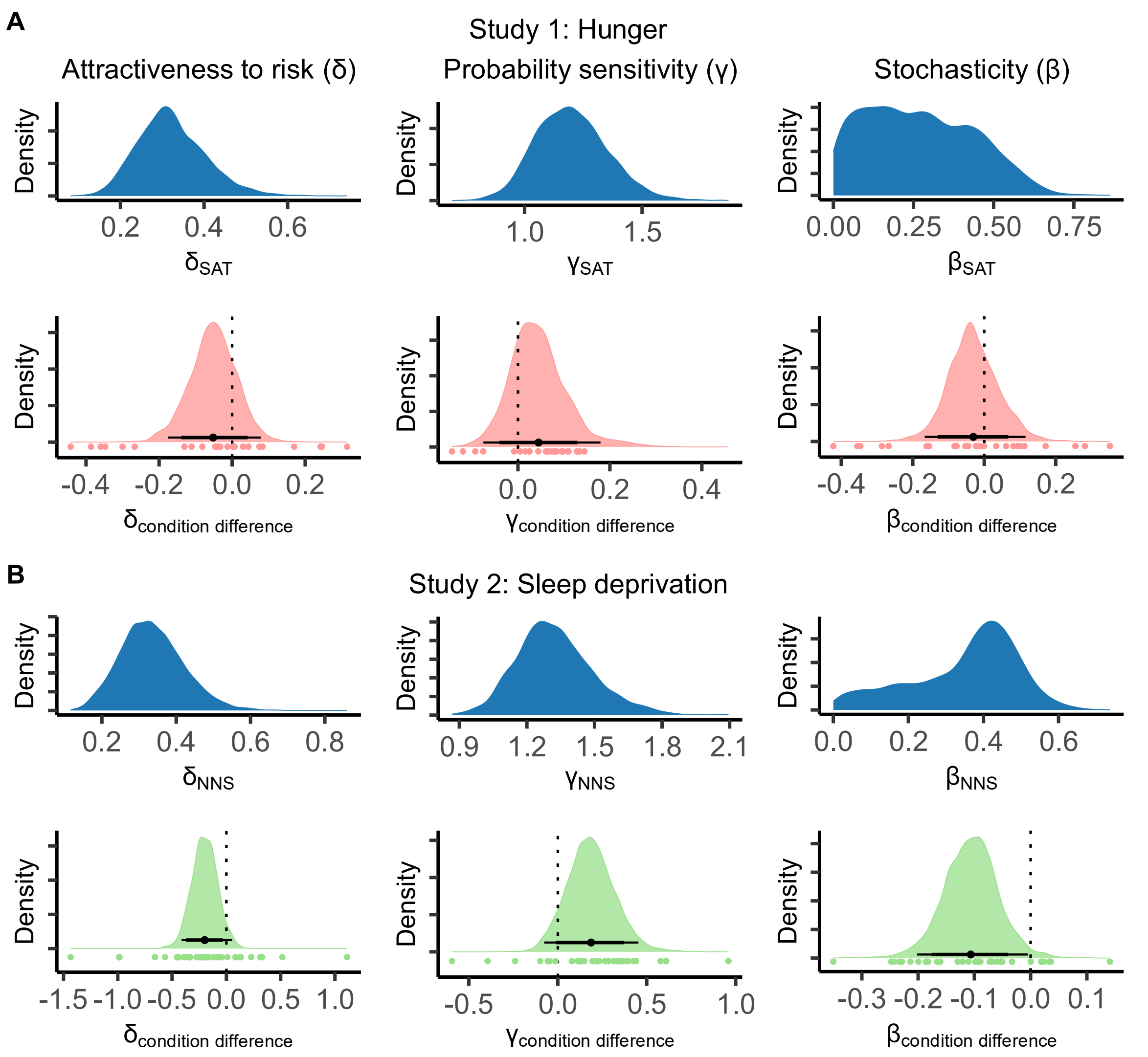


**Figure S5.** Posterior Distributions of Group-Level Parameters of the Prospect Theory Model with fMRI Participants. The posterior distributions of the group-level parameters for study 1 (hunger; A) and study 2 (sleep deprivation; B): attractiveness to risk $\delta$, sensitivity to probabilities $\gamma$, policy parameter $\beta$. Parameter values of the sated (A) and rested (B) condition are shown in blue, the difference between conditions is shown in red for study 1 and in green for study 2. The horizontal lines represent the highest distribution interval (HDI) of 95% (thin line) and 85% (thick line), with the mean of the distribution as dot. The coloured dots underneath the distribution represent the mean values of individual subjects. All parameters, in both studies, did not differ significantly between conditions as shown by the HDI containing zero in each difference distribution. Values are reported in Table S2 and Table S3.

***Prospect Theory with Ghrelin Coefficients***

Here (Table S4, Table S5, and Figure S6) we report that parameter values of the PT model with ghrelin coefficients fitted to the subset of participants of which the neuroimaging data was analysed. The values differ from those in Table 3 and Table 4 due to the different sample sizes. Additionally, this model contains the ghrelin coefficients for each of the PT parameters. The ghrelin coefficients are reported in the main article (Table 4 and Figure 6).

**Table S4.** Prospect Theory Parameter Values with Ghrelin Coefficients. Mean ($\pm$SD) of the blue distributions shown in Figure S6. Estimated parameters for the PT model with ghrelin coefficients for the baseline condition (i.e., sated or night of normal sleep) and the experimental condition (i.e., fasted or total sleep deprivation) at the group level. The model was fitted to the subset of participants of which the neuroimaging data was analysed. Attractiveness to risk $\delta$, sensitivity to probabilities $\gamma$, policy parameter $\beta$.

| | | Study 1 | Study 2 |
|---|---|---|---|
| $\delta$ | Baseline condition | $0.32\ (\pm 0.08)$ | $0.34\ (\pm 0.08)$ |
| | Experimental condition | $0.29\ (\pm 0.08)$ | $0.23\ (\pm 0.08)$ |
| $\gamma$ | Baseline condition | $1.21\ (\pm 0.15)$ | $1.33\ (\pm 0.17)$ |
| | Experimental condition | $1.27\ (\pm 0.16)$ | $1.6\ (\pm 0.22)$ |
| $\beta$ | Baseline condition | $0.27\ (\pm 0.17)$ | $0.35\ (\pm 0.15)$ |
| | Experimental condition | $0.24\ (\pm 0.19)$ | $0.24\ (\pm 0.15)$ |

**Table S5.** Conditional Differences of Prospect Theory with Ghrelin Coefficients. The difference between the baseline and experimental condition for each study (the coloured distributions in Figure S6).

| | Mean (±SD) | 95% HDI | $dBF_{10}$ |
|---|---|---|---|
| Study 1 | | | |
| Attractiveness to risk ($\delta$) | $-0.05$ ($\pm 0.06$) | $-0.17, 0.08$ | 0.31 |
| Sensitivity to probabilities ($\gamma$) | 0.05 ($\pm 0.07$) | $-0.08, 0.19$ | 3.49 |
| Policy parameter ($\beta$) | $-0.03$ ($\pm 0.08$) | $-0.19, 0.12$ | 0.48 |
| Study 2 | | | |
| Attractiveness to risk ($\delta$) | $-0.22$ ($\pm 0.12$) | $-0.47, 0.02$ | 0.04 |
| Sensitivity to probabilities ($\gamma$) | 0.24 ($\pm 0.14$) | $-0.04, 0.52$ | 24.55 |
| Policy parameter ($\beta$) | $-0.11$ ($\pm 0.05$) | $-0.21, -0.01$ | 0.02 |

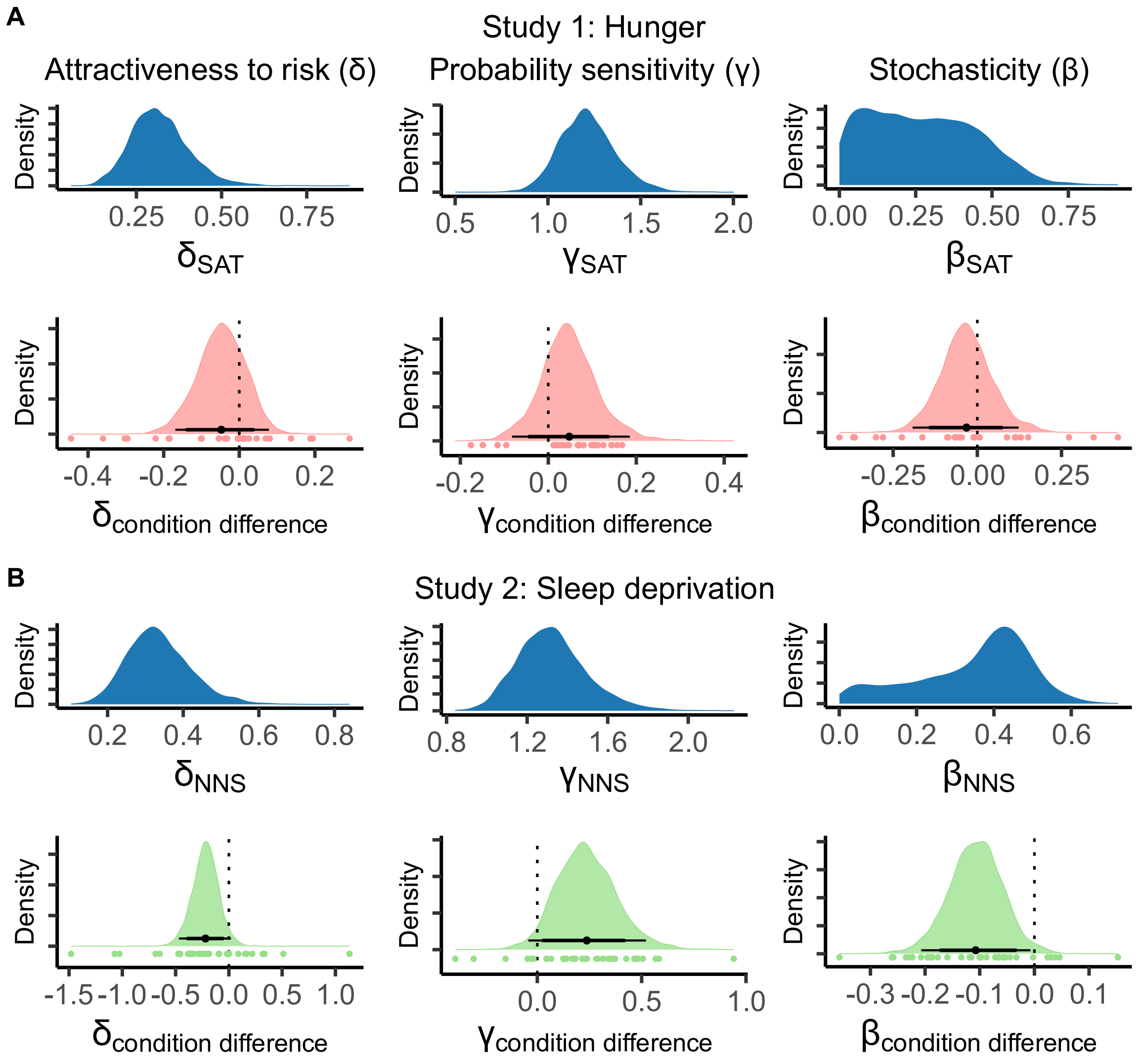


**Figure S6.** Posterior Distributions of Group-Level Parameters of the Prospect Theory Model with Ghrelin Coefficients. The posterior distributions of the group-level parameters for study 1 (hunger; A) and study 2 (sleep deprivation; B): attractiveness to risk $\delta$, sensitivity to probabilities $\gamma$, policy parameter $\beta$. Parameter values of the sated (A) and rested (B) condition are shown in blue, the difference between conditions is shown in red for study 1 and in green for study 2. The horizontal lines represent the highest distribution interval (HDI) of 95% (thin line) and 85% (thick line), with the mean of the distribution as dot. The coloured dots underneath the distribution represent the mean values of individual subjects. Most parameters, in both studies, did not differ significantly between conditions as shown by the HDI containing zero in each difference distribution. The exception is the stochasticity in study 2. Values are reported in Table S4 and Table S5.

**Neuroimaging Results**

***Subjective Value***

**Table S6.** Main Effect of Subjective Value Across Conditions. Regions with peak activity after small volumes corrections with the Rangel mask.

| Region | MNI coordinates | | | Peak T-value | $p(FWE)$ |
|---|---|---|---|---|---|
| | X | Y | Z | | |
| Study 1 | | | | | |
| Right posterior cingulate cortex | 2 | −34 | 38 | 7.48 | 0.000 |
| Right ventral striatum | 12 | 11 | −14 | 6.37 | 0.001 |
| Right anterior cingulate cortex | 8 | 42 | 10 | 6.3 | 0.001 |
| Left anterior cingulate cortex | −3 | 42 | 5 | 5.99 | 0.002 |
| Left nucleus caudatus | −12 | 12 | 2 | 4.86 | 0.027 |
| Study 2 | | | | | |
| Left nucleus caudatus | −9 | 11 | 4 | 5.43 | 0.003 |
| Left anterior cingulate cortex | −3 | 40 | 8 | 4.91 | 0.01 |
| Right posterior cingulate cortex | 2 | −32 | 34 | 4.53 | 0.027 |

***Choice***

**Table S7.** Main Effect of Risky Choice versus Safe Choice. Regions with peak activity of choice after small volumes corrections with the not-thresholded choice mask from Cui and colleagues (2022).

| Region | MNI coordinates | | | Peak T-value | $p(FWE)$ |
|---|---|---|---|---|---|
| | X | Y | Z | | |
| Study 1 | | | | | |
| Right dorsolateral prefrontal cortex | 50 | 42 | 18 | 7.41 | 0.000 |
| Right inferior parietal gyrus | 50 | −38 | 52 | 7.17 | 0.000 |
| Right inferior temporal gyrus | 60 | −44 | −14 | 7.09 | 0.000 |
| Left anterior insula | −32 | 22 | 2 | 6.92 | 0.000 |
| Right posterior cingulate cortex | 0 | −20 | 28 | 5.83 | 0.01 |
| Study 2 | | | | | |
| Left nucleus caudatus | −10 | 11 | 0 | 5.52 | 0.016 |
| Left dorsolateral prefrontal cortex | −40 | 53 | −2 | 5.31 | 0.035 |

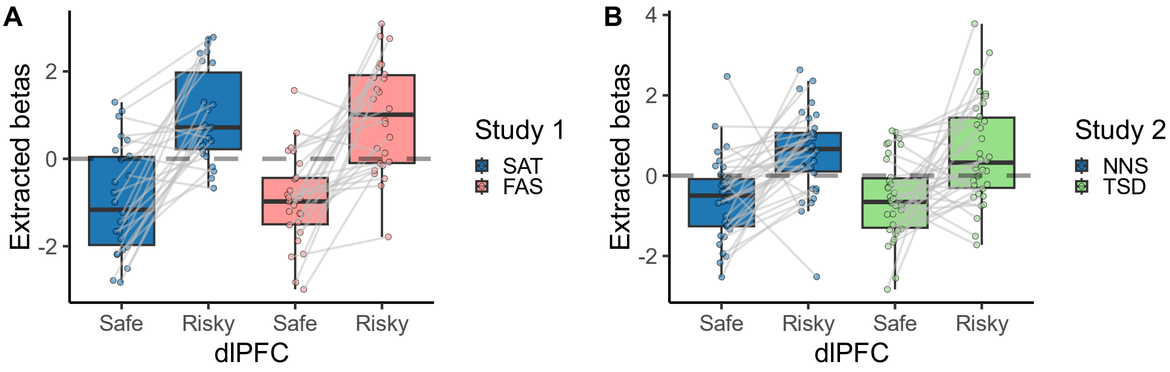


**Figure S7.** Extracted β-Coefficients of Choice. The extracted $\beta$-coefficients of each participant for the peak activity in the right dlPFC (study 1, A; $X = 50, Y = 42, Z = 18$) and the left dlPFC (study 2, B; $X = -40, Y = 53, Z = -2$). While there was an effect of choice in these regions, there was no interaction with condition.

***Ghrelin Correlation***

Here we report supporting evidence for the lack of effect of ghrelin on the neural activity of subjective value and choice. We extracted the $\beta$-coefficients from the different GLMs (subjective value and choice) for the brain regions with significant activity and where we initially predicted to be activity related to either subjective value or choice. For the subjective value model, this was the PCC and ventral striatum in study 1, and the PCC in study 2. For the choice model, this was the dlPFC in both studies (see Table S6 and Table S7 for the MNI coordinates of the relevant regions for subjective value and choice respectively). The difference in $\beta$-coefficients between the experimental condition (fasted or sleep deprived) and the baseline condition (sated or normal sleep) is plotted against the difference in ghrelin levels between conditions. None of the correlations were statistically significant (see Table S8 and Figure S8).

**Table S8.** Correlation Between Ghrelin Difference and $\beta$ Difference. Pearson correlation coefficient (p-values) of the difference in ghrelin levels between conditions, and for subjective value the difference in $\beta$-coefficients between the experimental condition (fasted or sleep deprived) and the baseline condition (sated or normal sleep), for choice it is the difference in $\beta$-coefficients between the risky and safe choice.

| | Study 1 | Study 2 |
|---|---|---|
| Subjective value | | |
| Posterior cingulate cortex | $-0.35\ (.08)$ | $0.15\ (.4)$ |
| Ventral striatum | $-0.24\ (.25)$ | – |
| Choice | | |
| Dorsolateral prefrontal cortex | $0.37\ (.07)$ | $0.18\ (.32)$ |

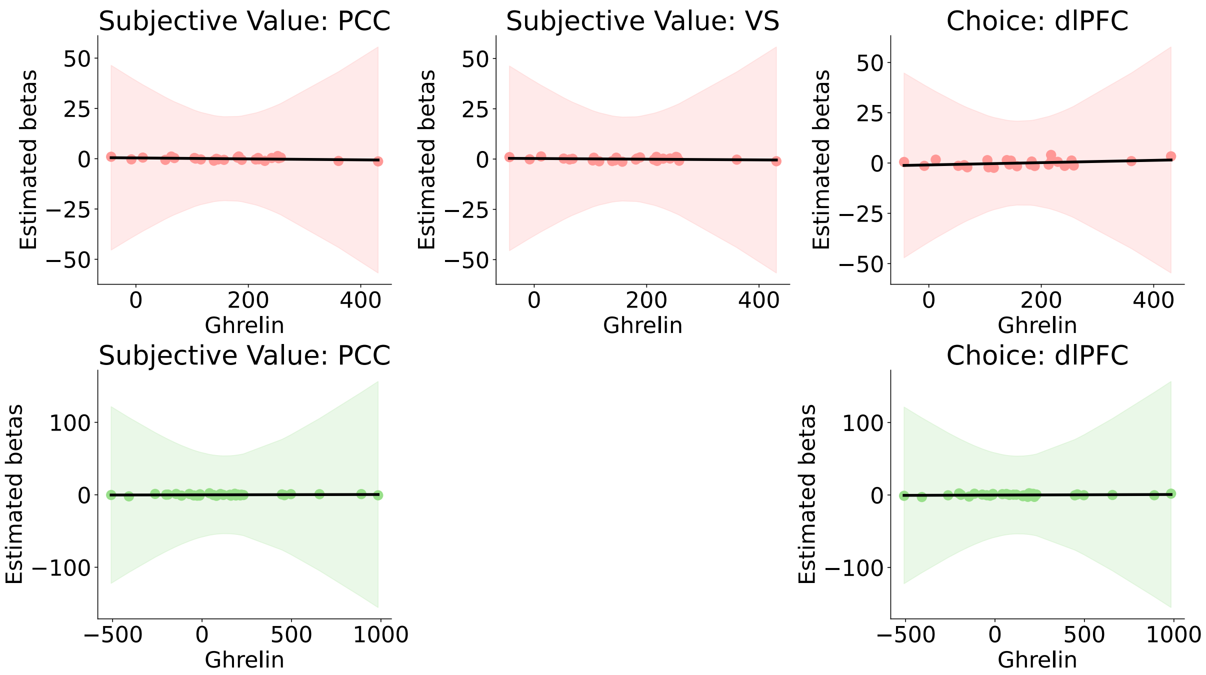


**Figure S8.** Estimated $\beta$-Coefficients of fMRI Models. The difference in extracted $\beta$-coefficients between conditions of the subjective value and choice GLMs for the neuroimaging data and their relationship to the difference in ghrelin levels between conditions. The black line is the best fitting regression line with coloured confidence bounds. The correlation was never statistically significant. Study 1 (fasting) in red, study 2 (sleep deprivation) in green; posterior cingulate cortex (PCC), ventral striatum (VS), dorsolateral prefrontal cortex (dlPFC).

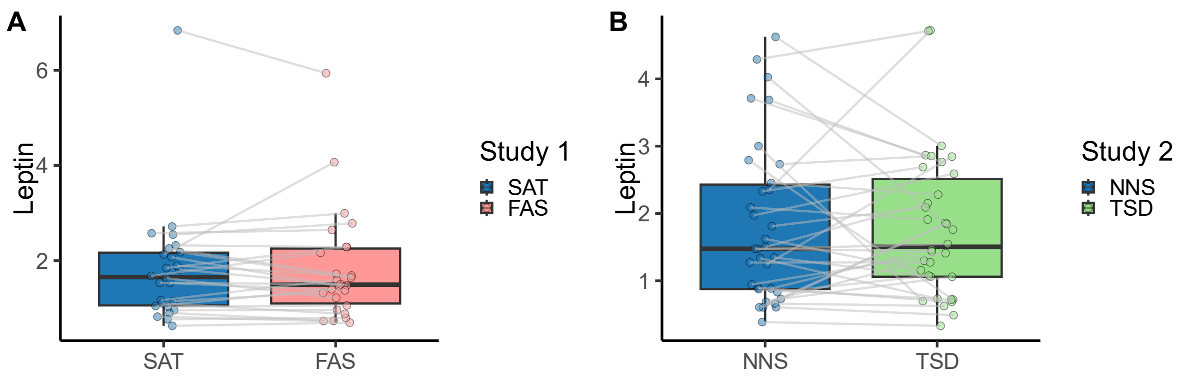


**Figure S9.** Leptin Levels. The data provided strong evidence against an increase in leptin levels between sated (A, blue; $Mdn_{SAT} = 1.66$) and the fasted condition (A, red; $Mdn_{FAS} = 1.5, BF = 0.21 \pm 0.03\%$) in study 1. Similarly, in study 2, the data provided strong evidence against an increase in leptin levels after one night of total sleep deprivation (B; $Mdn_{NNS} = 1.48, Mdn_{TSD} = 1.51, BF = 0.19 \pm 0.04\%$).

**Controlling for Insulin, Cortisol, Leptin, and Glucose**

Ghrelin is not the only food related hormone that could have an effect on risky decision-making. Insulin and leptin interact with dopaminergic reward system as well. Therefore, we controlled for their effects when analysing the effects of ghrelin, the main focus of the article, on risky decision-making. Before controlling for the other endocrine variables, we investigated their correlations. This was done for the difference between the conditions as well as for the values in each condition separately. The difference between both conditions in study 1 (Figure S10) in ghrelin had a large negative correlation with insulin $(r = -0.66, p < 0.001)$. In addition, there was a significant positive correlation between insulin and glucose $(r = 0.67, p < 0.001)$. The comparison of the endocrine levels in either the sated (Figure S11) or fasted (Figure S12) condition revealed similar results. The differences between conditions in study 2 (Figure S13) in insulin correlated with leptin $(r = 0.43, p < 0.05)$. This correlation was bigger in after one night of normal sleep ($r = 0.65, p < 0.001$; Figure S14) but lost its significance after one night of total sleep deprivation ($r = 0.23$; Figure S15).

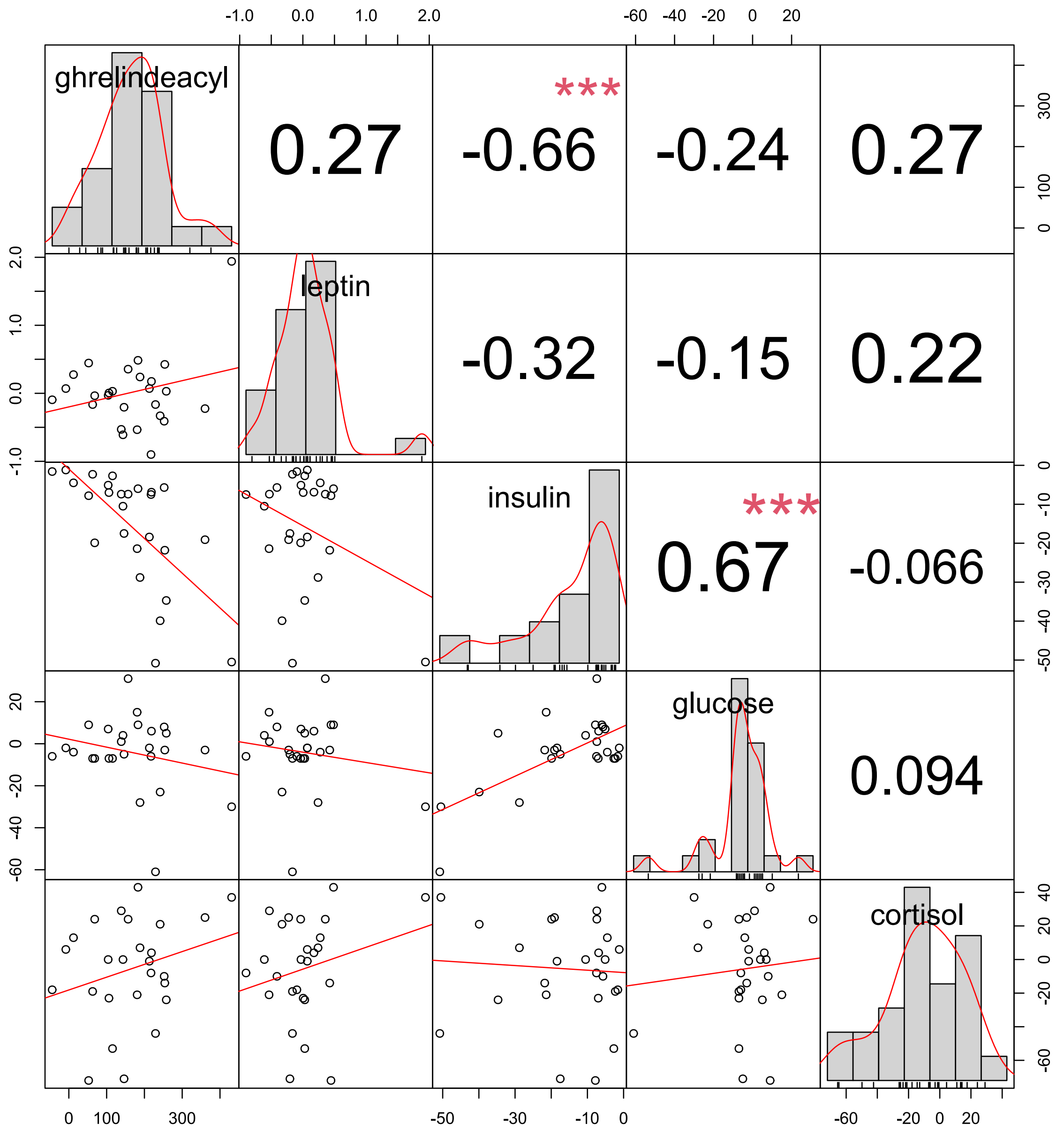


**Figure S10.** Overview of Correlations for Condition Differences in Study 1. The diagonal contains histograms of the difference between fasted and sated for the different endocrine variables. The scatterplots below the diagonal show the best fitting regression in red. Above the diagonal are the Pearson correlations with stars representing their significance $(0 < *** < 0.001 < ** < 0.01 < * < 0.05 < . < 0.1)$.

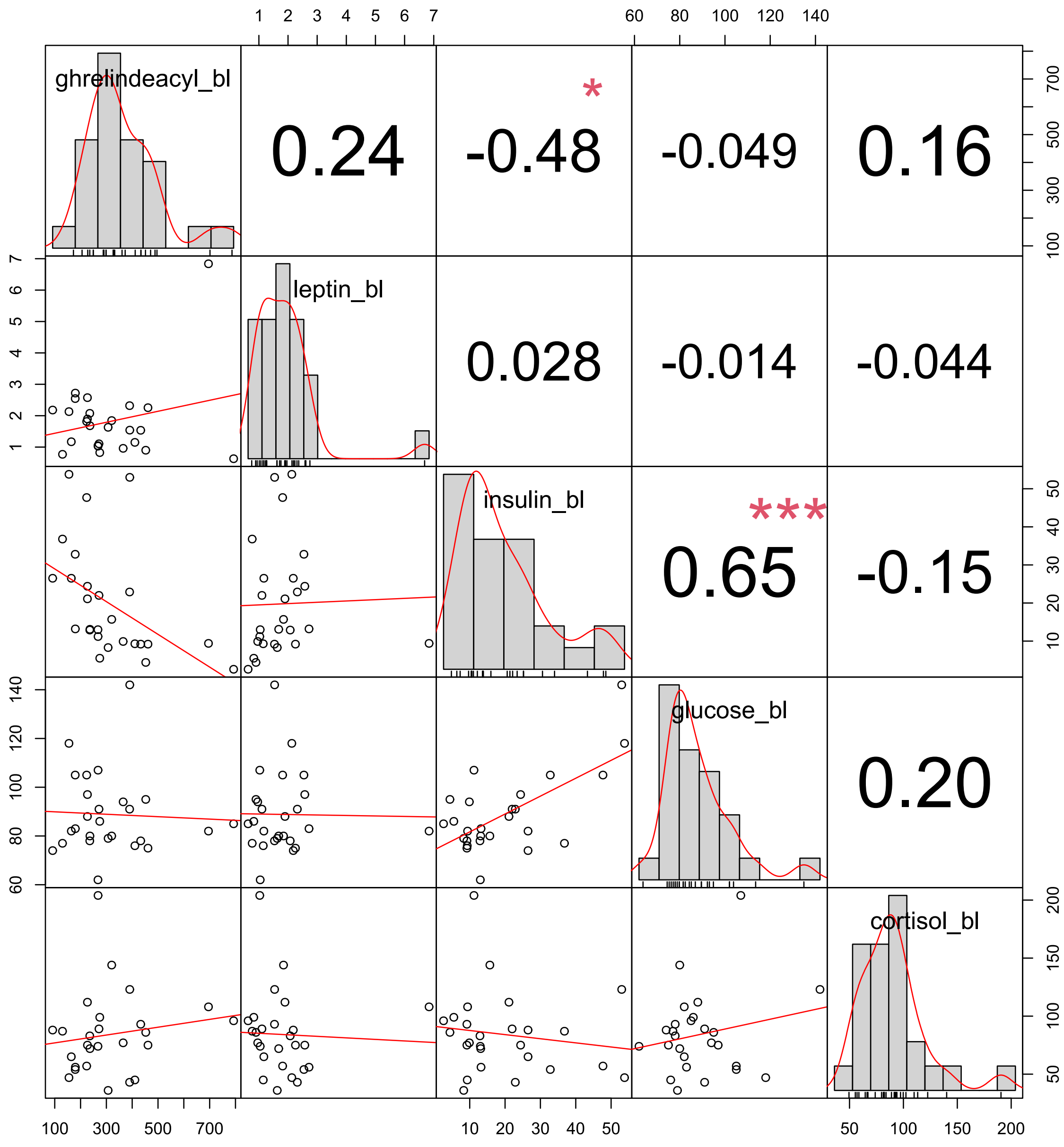


**Figure S11.** Overview of Correlations in the Baseline Condition of Study 1. The diagonal contains histograms of the different endocrine variables when sated. The scatterplots below the diagonal show the best fitting regression in red. Above the diagonal are the Pearson correlations with stars representing their significance ($0 < *** < 0.001 < ** < 0.01 < * < 0.05 < . < 0.1$).

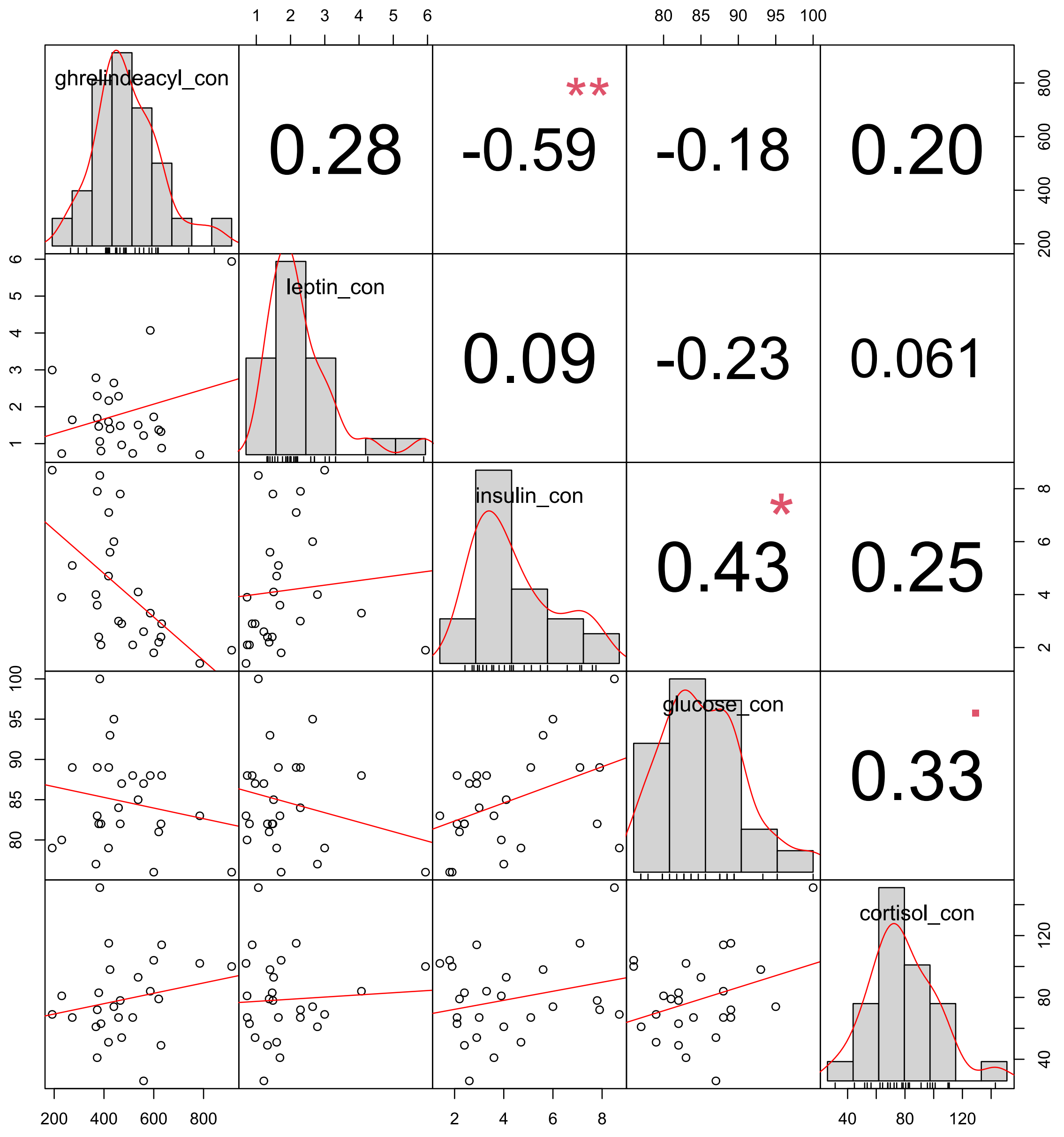


**Figure S12.** Overview of Correlations in the Experimental Condition of Study 1. The diagonal contains histograms of the different endocrine variables when fasted. The scatterplots below the diagonal show the best fitting regression in red. Above the diagonal are the Pearson correlations with stars representing their significance ($0 < *** < 0.001 < ** < 0.01 < * < 0.05 < . < 0.1$).

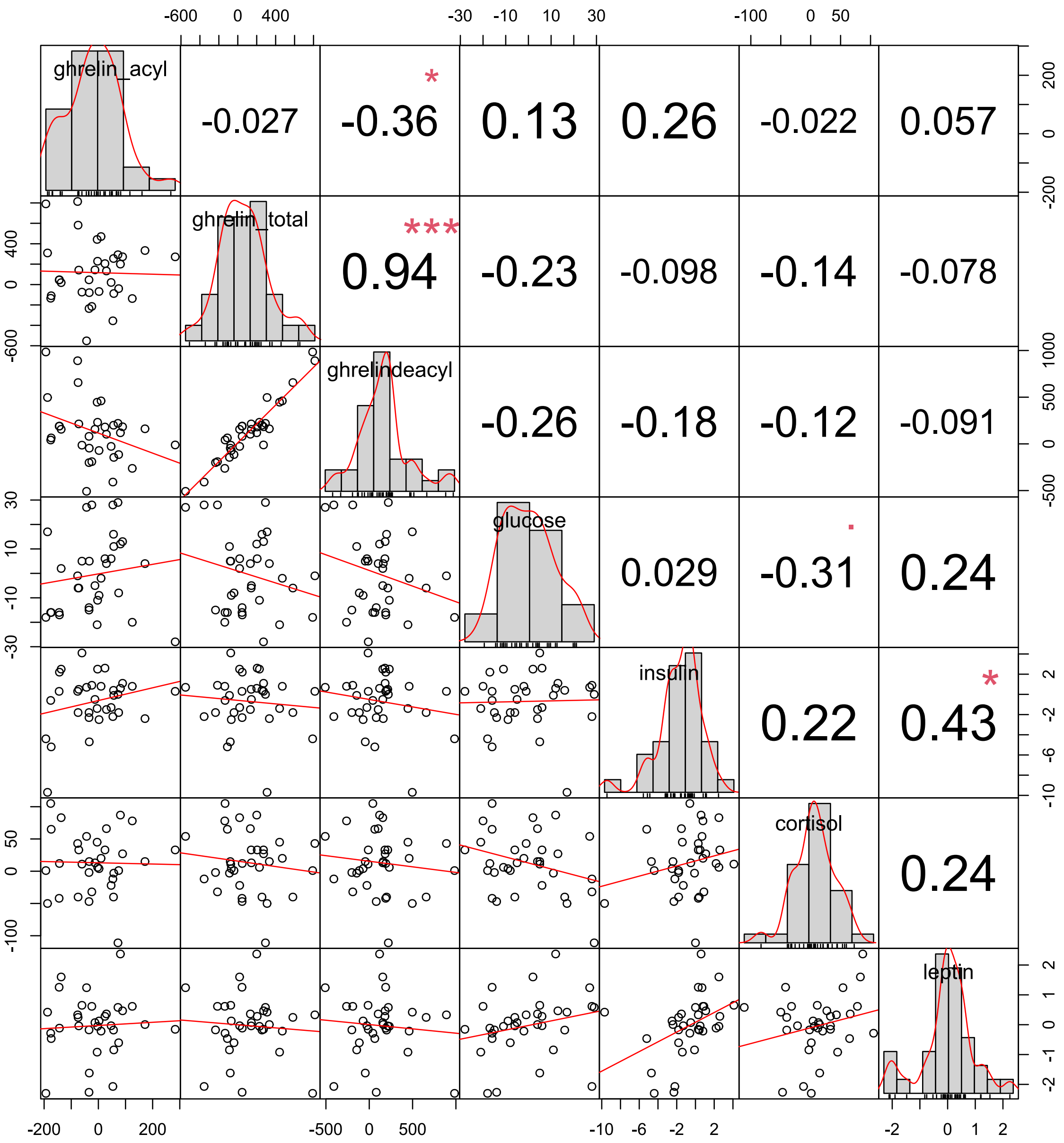


**Figure S13.** Overview of Correlations for Condition Differences in Study 2. The diagonal contains histograms of the difference between one night of total sleep deprivation and one night of normal sleep for the different endocrine variables. The scatterplots below the diagonal show the best fitting regression in red. Above the diagonal are the Pearson correlations with stars representing their significance ($0 < *** < 0.001 < ** < 0.01 < * < 0.05 < . < 0.1$).

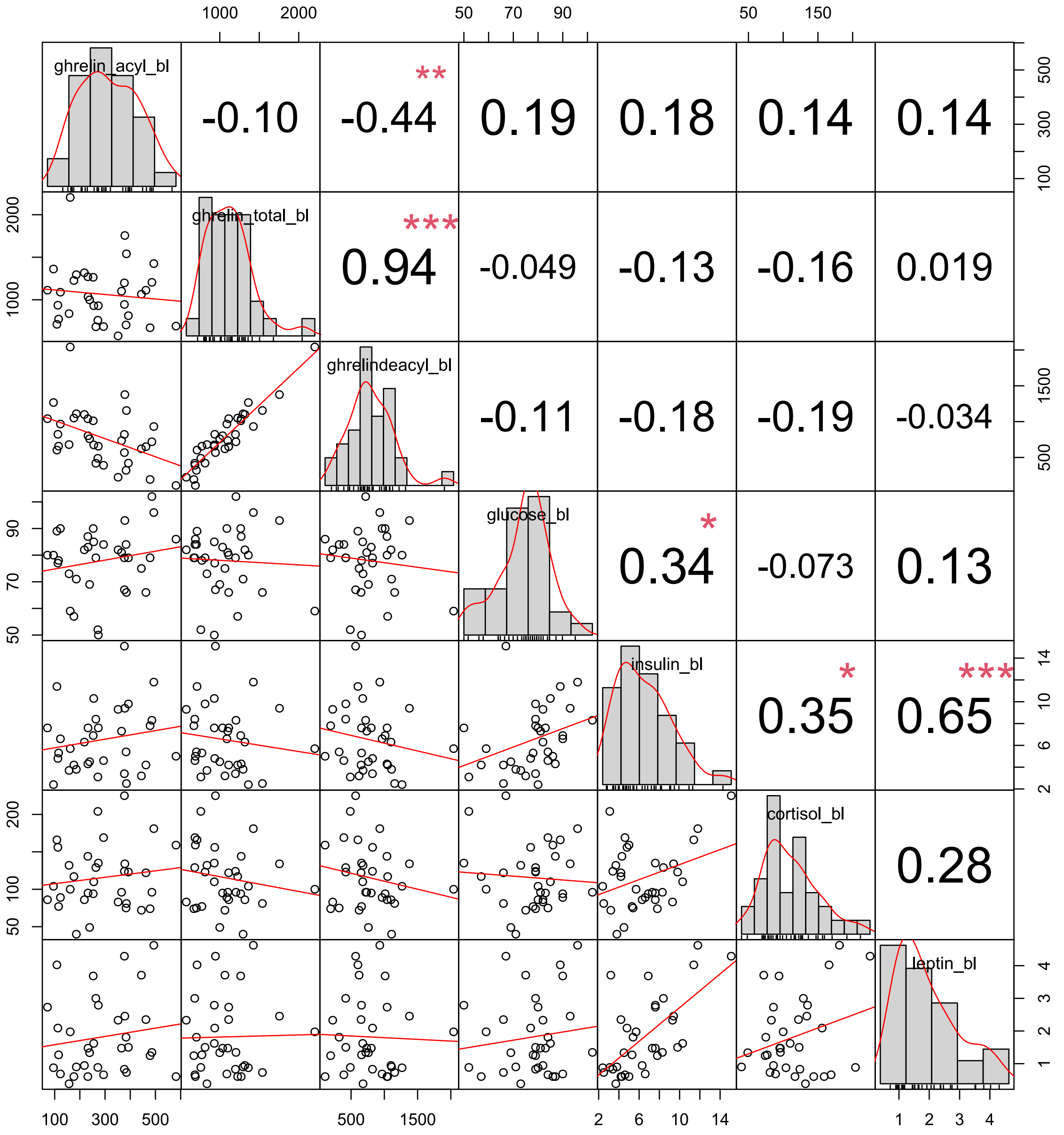


**Figure S14.** Overview of Correlations in the Baseline Condition of Study 2. The diagonal contains histograms of the different endocrine variables after one night of normal sleep. The scatterplots below the diagonal show the best fitting regression in red. Above the diagonal are the Pearson correlations with stars representing their significance ($0 < *** < 0.001 < ** < 0.01 < * < 0.05 < . < 0.1$).

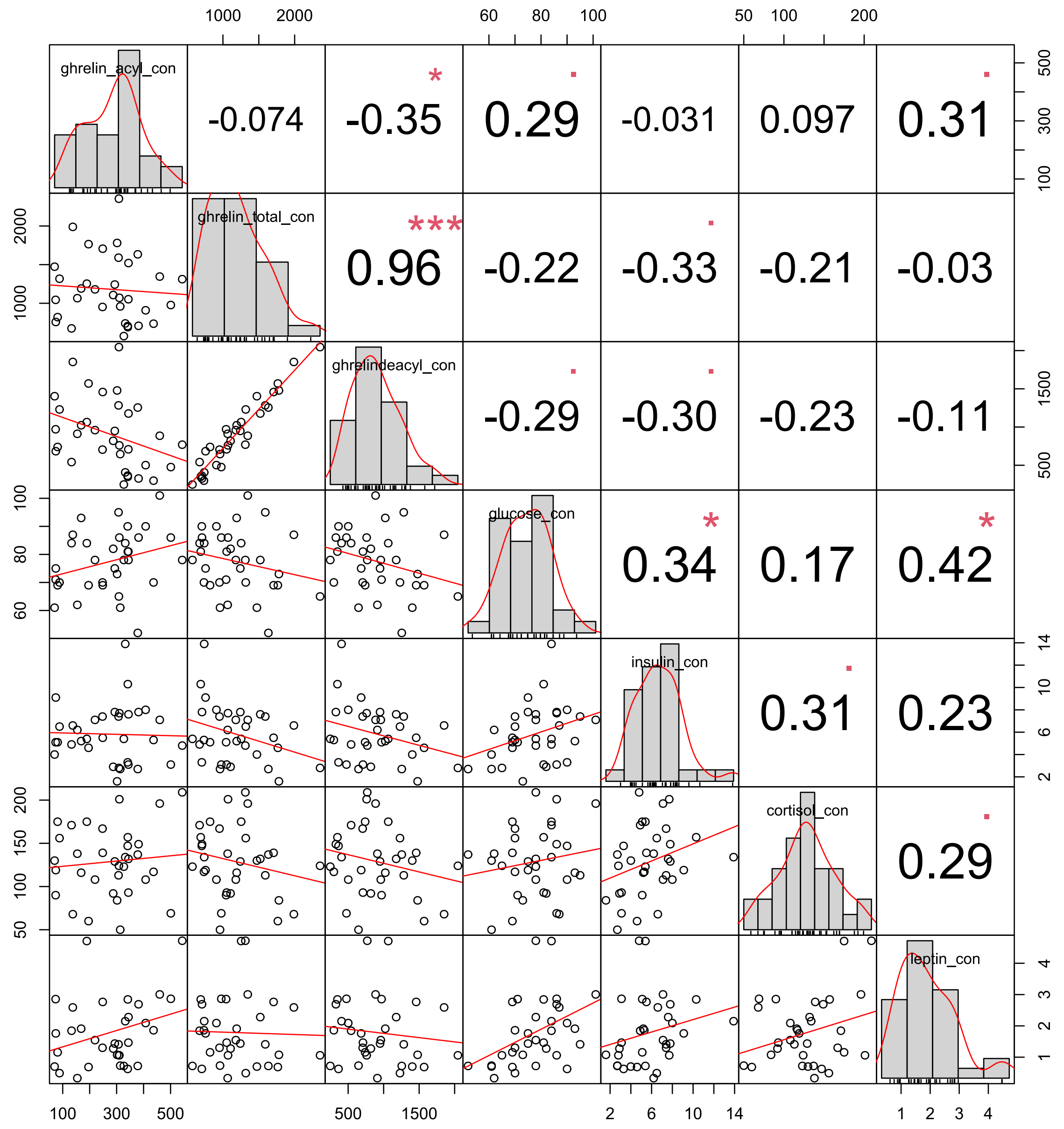


**Figure S15.** Overview of Correlations in the Experimental Condition of Study 2. The diagonal contains histograms of the different endocrine variables after one night of total sleep deprivation. The scatterplots below the diagonal show the best fitting regression in red. Above the diagonal are the Pearson correlations with stars representing their significance $(0 < *** < 0.001 < ** < 0.01 < * < 0.05 < . < 0.1)$.

***Insulin***

Insulin is a peptide that fluctuates with levels of food intake, and thus covaries with hunger. Contrary to ghrelin, there was strong evidence for a decrease in insulin levels in the fasting condition ($Mdn_{FAS} = 4.15$), compared to the sated condition ($Mdn_{SAT} = 19.78, BF = 1959.13 \pm 0\%$) in study 1 (Figure S16.A). In study 2, there was anecdotal evidence against a difference between insulin levels after one night of normal sleep ($Mdn_{NNS} = 6$) and insulin levels after one night of total sleep deprivation ($Mdn_{TDS} = 5.4, BF = 0.53 \pm 0.03\%$; Figure S16.B).

We reanalysed the difference in ghrelin levels between conditions, now by controlling for the insulin levels. For this, we compared a Bayesian linear model with insulin as a predictor for ghrelin to a model with insulin and the condition as predictive values. When controlled for insulin, there was anecdotal evidence for no difference in ghrelin between both conditions in study 1 ($BF = 0.85 \pm 1.6\%$) and study 2 ($BF = 0.4 \pm 1.19\%$). This suggests that the changes in ghrelin levels were better explained by the changes in insulin levels than by the experimental manipulation.

To control for the effect of insulin in the computational model, we included insulin parameters for each model parameter. Using $\delta$ as an example and building on Equation 9 and Equation 10, the model was adjusted with

$$\delta_e = \delta_g + \delta_i = \Delta_g * \lambda_{g\delta} + \Delta_i * \lambda_{i\delta} \tag{S1}$$

where $\delta_e$ is the sum of the ghrelin controlled parameter $\delta_g$ and the insulin controlled parameter $\delta_i$ which contains the difference in insulin between conditions $\Delta_i$, and the insulin coefficient $\lambda_{i\delta}$ for the model's parameter, in this example $\delta$. The same is done for $\gamma_e$ and $\beta_e$. This resulted in the a PT model where the subjective probability weight (Equation 2) is

$$w(P_t) = \frac{\left(\delta + c_t * (\delta_s + \delta_e)\right) * P_t^{\gamma + c_t * (\gamma_s + \gamma_e)}}{\left(\delta + c_t * (\delta_s + \delta_e)\right) * P_t^{\gamma + c_t * (\gamma_s + \gamma_e)} + (1 - P_t)^{\gamma + c_t * (\gamma_s + \gamma_e)}} \tag{2}$$

The model was fitted on the subset of fMRI participants. To verify that the model was able to describe the observed behaviour, we compared the simulated to the observed risky choice selection. There was a large overlap between the predicted and observed behaviour (Figure S17).

With the inclusion of ghrelin and insulin, the model parameters did not differ between conditions according to the $95\%$ HDI, except for $\beta$ in study 2 (Figure S18). Attractiveness to risk $\delta$ in study 2 showed a tendency towards significance but the zero was still captured by the $95\%$ HDI. The inclusion of the covariates changed the direction of the difference between conditions, when compared to the model without covariates. For example, in study 1, the choice stochasticity decreased when controlling for insulin, while it increased when only ghrelin was included. Comparing the effects of both covariates, it is noteworthy that ghrelin and leptin affect the different parameters in opposite directions, except for $\gamma$ and $\beta$ in study 2. Both increase the difference of $\gamma$ between conditions, and decrease the difference of $\beta$ between conditions.

The neural activity related to choice did not show an effect of ghrelin. However, when controlling for insulin, one region showed significant activity for the interaction effect of choice (risky versus safe) and condition (fasted versus sated) in study 1 (Table S13).

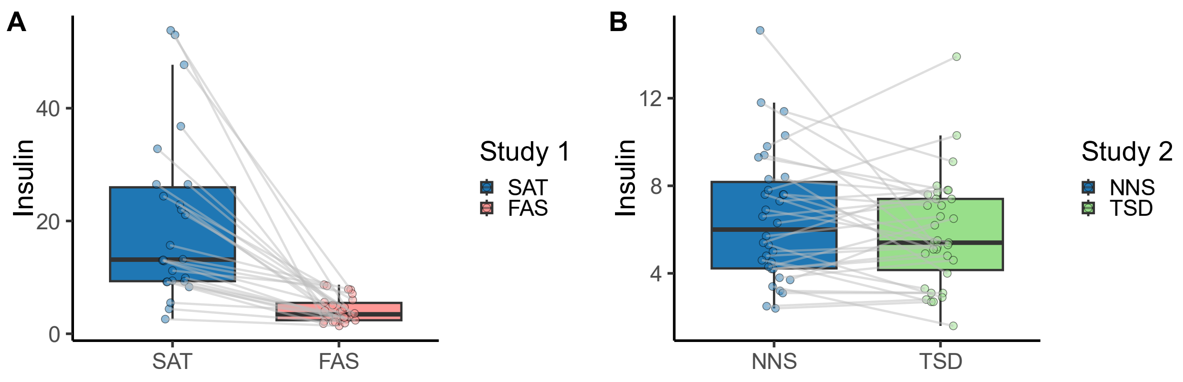


**Figure S16**. Insulin Levels. A) Insulin decreased in the fasted condition (red; $Mdn_{FAS} = 4.15$) compared to the sated condition (blue; $Mdn_{SAT} = 19.78$) in study 1. B) There were no changes in insulin levels between one night of normal sleep (blue; $Mdn_{NNS} = 6$) or one night of total sleep deprivation (green; $Mdn_{TSD} = 5.4$) in study 2.

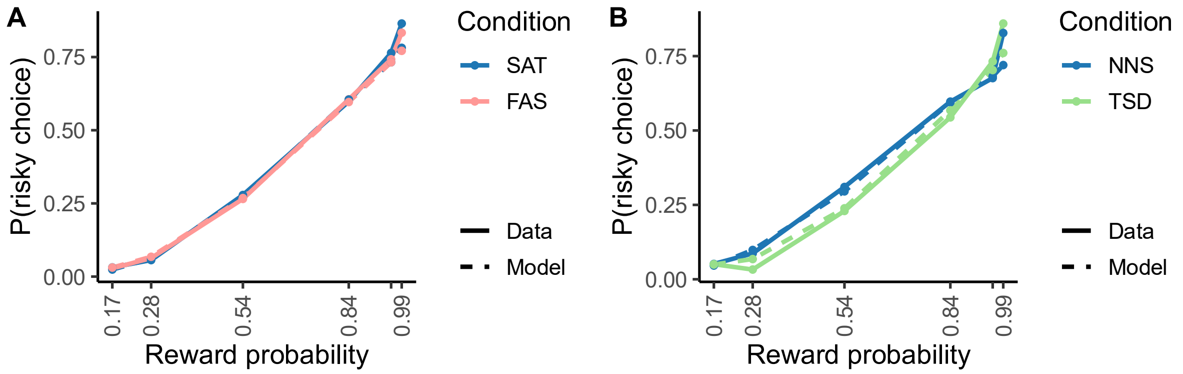


**Figure S17.** Choice Predictions and Behavioural Data of the Prospect Theory Model with Ghrelin and Insulin Covariates. The probability of selecting the risky option per reward probability $(0.17, 0.28, 0.54, 0.84, 0.96, 0.99)$ for participants (full line) and simulated by the prospect theory model (dashed line). The model predictions show a large overlap with the behavioural data.

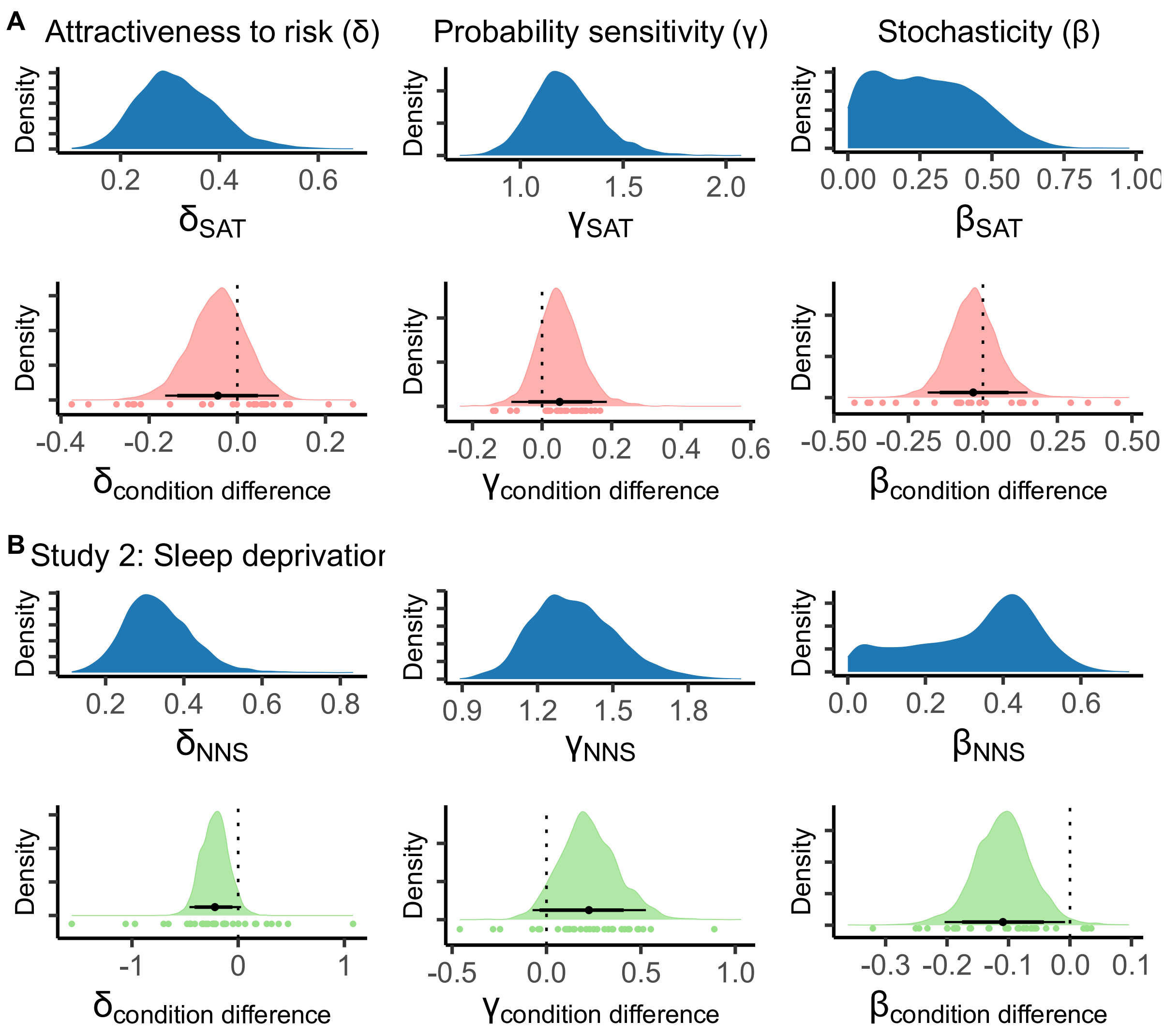


**Figure S18.** Posterior Distribution of Group-Level Parameters of the Prospect Theory Model with Ghrelin and Insulin Covariates.

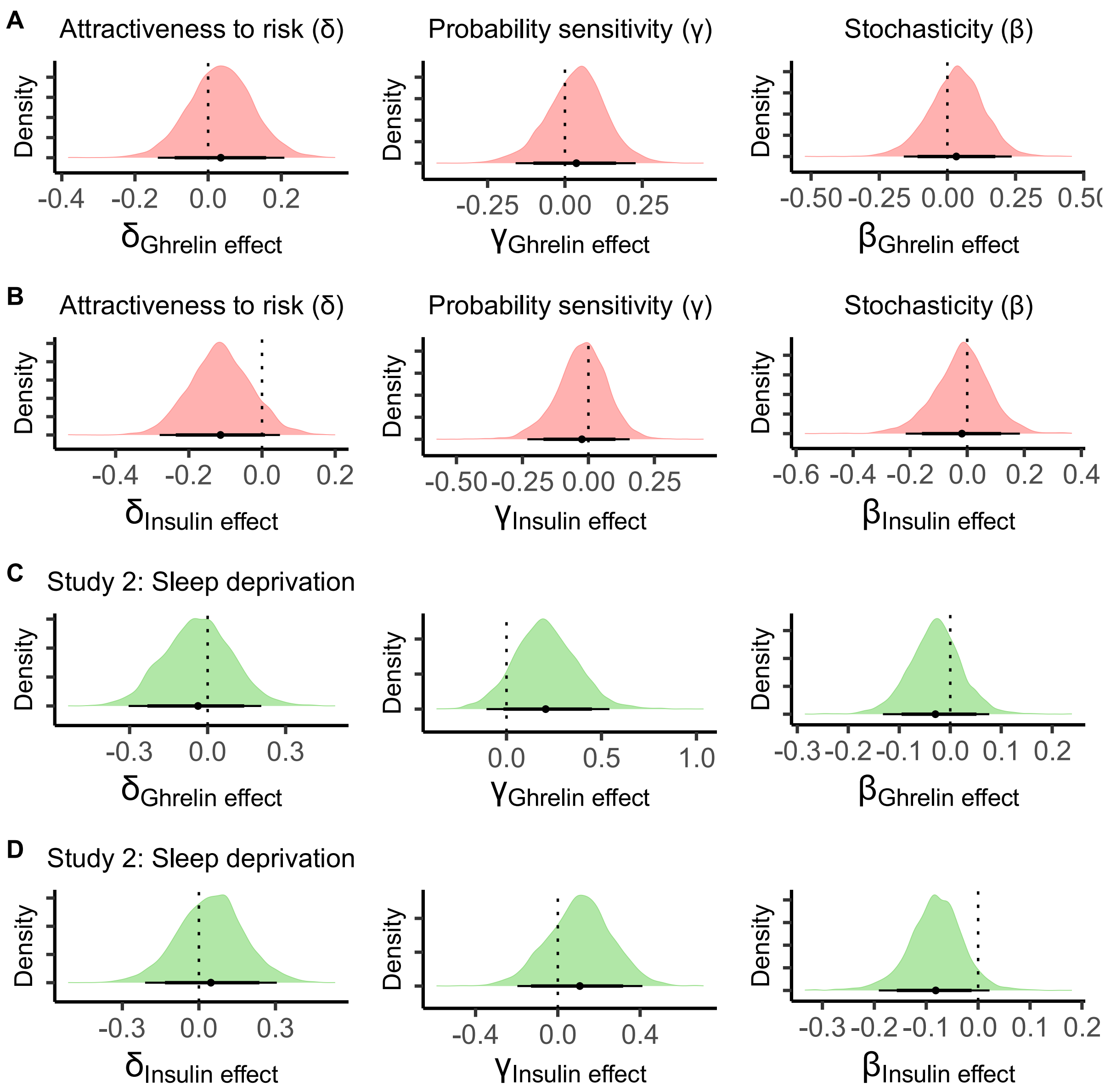


**Figure S19.** Posterior Distribution of Ghrelin Coefficients for the Parameters of the Prospect Theory Model. The posterior distributions of the ghrelin and leptin coefficients for the group-level parameters for study 1 (hunger; A and B) and study 2 (sleep deprivation; C and D): attractiveness to risk δ, sensitivity to probabilities γ, policy parameter β. Zero falls within the highest density intervals (HDI; thick bars: 0.85, thin bars: 0.95) for all parameter in both studies.

**Table S9.** Parameter Values of Prospect Theory with Ghrelin and Insulin Covariates. The mean (±SD) of the blue distributions shown in Figure S18: the estimated parameters for the PT model for the baseline condition (i.e., sated or night of normal sleep) and the experimental condition (i.e., fasted or total sleep deprivation) at the group level. Attractiveness to risk $\delta$, sensitivity to probabilities $\gamma$, policy parameter $\beta$.

| | | Study 1 | Study 2 |
|---|---|---|---|
| $\delta$ | Baseline condition | $0.32\ (\pm 0.08)$ | $0.33\ (\pm 0.09)$ |
| | Experimental condition | $0.29\ (\pm 0.08)$ | $0.23\ (\pm 0.08)$ |
| $\gamma$ | Baseline condition | $1.22\ (\pm 0.17)$ | $1.35\ (\pm 0.17)$ |
| | Experimental condition | $1.28\ (\pm 0.18)$ | $1.61\ (\pm 0.22)$ |
| $\beta$ | Baseline condition | $0.27\ (\pm 0.17)$ | $0.33\ (\pm 0.14)$ |
| | Experimental condition | $0.24\ (\pm 0.19)$ | $0.22\ (\pm 0.16)$ |

**Table S10.** Condition Differences for the Prospect Theory With Ghrelin and Insulin Covariates. The difference in parameter values between the baseline and experimental condition for each study (the coloured distributions in Figure S18).

| | Mean (±SD) | 95% HDI | $dBF_{10}$ |
|---|---|---|---|
| Study 1 | | | |
| Attractiveness to risk ($\delta$) | $-0.04\ (\pm 0.06)$ | $-0.16, 0.09$ | $0.33$ |
| Sensitivity to probabilities ($\gamma$) | $0.05\ (\pm 0.07)$ | $-0.09, 0.19$ | $3.63$ |
| Policy parameter ($\beta$) | $-0.03\ (\pm 0.08)$ | $-0.19, 0.15$ | $0.49$ |
| Study 2 | | | |
| Attractiveness to risk ($\delta$) | $-0.22\ (\pm 0.13)$ | $-0.46, 0.03$ | $0.05$ |
| Sensitivity to probabilities ($\gamma$) | $0.22\ (\pm 0.15)$ | $-0.07, 0.53$ | $14.04$ |
| Policy parameter ($\beta$) | $-0.11\ (\pm 0.05)$ | $-0.2, -0.01$ | $0.02$ |

**Table S11.** Ghrelin Coefficients of the Prospect Theory Parameters. The effect of ghrelin on the estimated parameter values of the PT model presented in Figure S19 (A and C).

| | Mean (±SD) | 95% HDI | $dBF_{10}$ |
|---|---|---|---|
| Study 1 | | | |
| Attractiveness to risk ($\delta$) | $0.03\ (\pm 0.09)$ | $-0.14, 0.21$ | $1.91$ |
| Sensitivity to probabilities ($\gamma$) | $0.04\ (\pm 0.1)$ | $-0.16, 0.23$ | $2.01$ |
| Policy parameter ($\beta$) | $0.03\ (\pm 0.1)$ | $-0.15, 0.24$ | $1.79$ |
| Study 2 | | | |
| Attractiveness to risk ($\delta$) | $-0.04\ (\pm 0.13)$ | $-0.3, 0.21$ | $0.64$ |
| Sensitivity to probabilities ($\gamma$) | $0.21\ (\pm 0.17)$ | $-0.11, 0.54$ | $8.87$ |
| Policy parameter ($\beta$) | $-0.03\ (\pm 0.05)$ | $-0.13, -0.08$ | $0.38$ |

**Table S12.** Insulin Coefficients of the Prospect Theory Parameters. The effect of insulin on the estimated parameter values of the PT model presented in Figure S19 (B and D).

| | Mean (±SD) | 95% HDI | $dBF_{10}$ |
|---|---|---|---|
| Study 1 | | | |
| Attractiveness to risk ($\delta$) | $-0.11\ (\pm 0.08)$ | $-0.28, 0.05$ | $0.1$ |
| Sensitivity to probabilities ($\gamma$) | $-0.02\ (\pm 0.1)$ | $-0.23, 0.16$ | $0.7$ |
| Policy parameter ($\beta$) | $-0.02\ (\pm 0.1)$ | $-0.22, 0.18$ | $0.76$ |
| Study 2 | | | |
| Attractiveness to risk ($\delta$) | $0.05\ (\pm 0.13)$ | $-0.21, 0.31$ | $1.82$ |
| Sensitivity to probabilities ($\gamma$) | $0.11\ (\pm 0.16)$ | $-0.2, 0.41$ | $3.11$ |
| Policy parameter ($\beta$) | $-0.08\ (\pm 0.05)$ | $-0.19, -0.02$ | $0.06$ |

**Table S13.** Interaction Effect of Choice and Condition. Regions with peak activity for the interaction of choice (risky versus safe) and condition (fasted versus sated) after small volumes corrections with the not-thresholded choice mask from Cui and colleagues (2022).

| Region | MNI coordinates | | | Peak T-value | $p(FWE)$ |
|---|---|---|---|---|---|
| | X | Y | Z | | |
| Study 1 | | | | | |
| Right dorsolateral prefrontal cortex | 22 | 2 | 46 | 5.61 | 0.000 |

***Cortisol***

Cortisol is a hormone that fluctuates with levels of food intake, and thus covaries with hunger. Contrary to ghrelin, there was moderate evidence against a difference in cortisol levels between the sated ($Mdn_{SAT} = 80$) and fasted condition ($Mdn_{FAS} = 76,\ BF = 0.31 \pm 0.03\%$) in study 1 (Figure S20.A). In study 2, there was anecdotal evidence against a difference between cortisol levels after one night of normal sleep ($Mdn_{NNS} = 107$) and cortisol levels after one night of total sleep deprivation ($Mdn_{TDS} = 129.5, BF = 0.64 \pm 0.03\%$; Figure S20.B).

We reanalysed the difference in ghrelin levels between conditions, now by controlling for the cortisol levels. For this, we compared a Bayesian linear model with cortisol as a predictor for ghrelin to a model with cortisol and the condition as predictive values. When controlled for cortisol, there was very strong evidence in favour of a difference in ghrelin levels between conditions in study 1 ($BF = 67.9 \pm 0.77\%$). However, there was anecdotal evidence against a difference in ghrelin between conditions in study 2 ($BF = 0.65 \pm 1.16\%$).

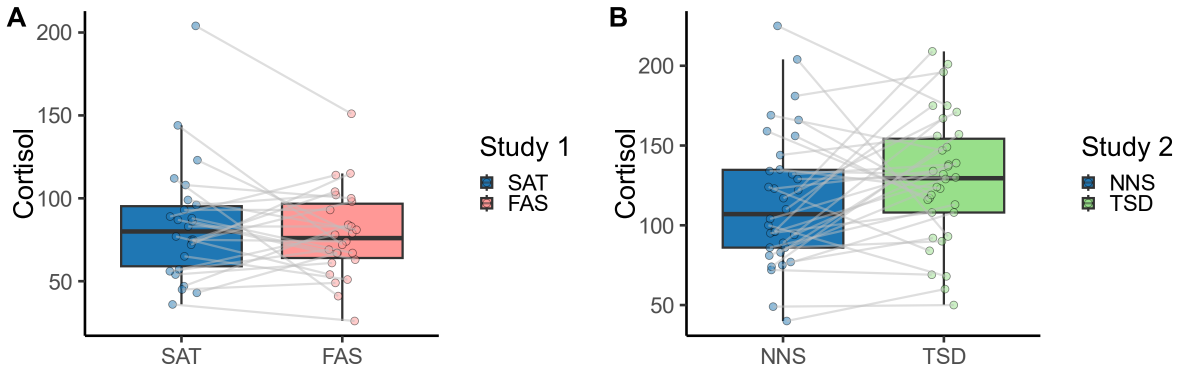


**Figure S20.** Cortisol Levels. A) Cortisol did not differ between the sated ($Mdn_{SAT} = 80$) and fasted condition (red; $Mdn_{FAS} = 76$) in study 1. B) There were no changes in cortisol levels between one night of normal sleep (blue; $Mdn_{NNS} = 107$) or one night of total sleep deprivation (green; $Mdn_{TSD} = 129.5$) in study 2.

***Leptin***

Leptin is a hormone that fluctuates with levels of food intake, and thus covaries with hunger. Contrary to ghrelin, there was strong evidence against differences in leptin between the sated ($Mdn_{SAT} = 1.66$) and fasted ($Mdn_{FAS} = 1.5, BF = 0.21 \pm 0.03\%$) condition in study 1 (Figure S9.A). Similarly, the data in study 2 (Figure S9.B) provided strong evidence against differences in leptin levels after one night of normal sleep ($Mdn_{NNS} = 1.48$) and leptin levels after one night of total sleep deprivation ($Mdn_{TSD} = 1.51, BF = 0.19 \pm 0.04\%$).

We reanalysed the difference in ghrelin levels between conditions, now by controlling for the leptin levels. For this, we compared a Bayesian linear model with leptin as a predictor for ghrelin to a model with leptin and the condition as predictive values. When controlled for leptin, there was very strong evidence in favour of a difference in ghrelin levels between conditions in study 1 ($BF = 59.41 \pm 1.51\%$). However, there was anecdotal evidence against a difference in ghrelin between conditions in study 2 ($BF = 0.5 \pm 0.96\%$).

***Glucose***

Glucose is a hormone that fluctuates with levels of food intake, and thus covaries with hunger. Contrary to ghrelin, there was moderate evidence against a difference in glucose levels between the sated ($Mdn_{SAT} = 84$) and fasted condition ($Mdn_{FAS} = 83.5,\ BF = 0.4 \pm 0.03\%$) in study 1 (Figure 21.A). In study 2, there was moderate evidence against a difference between glucose levels after one night of normal sleep ($Mdn_{NNS} = 79.5$) and glucose levels after one night of total sleep deprivation ($Mdn_{TDS} = 78, BF = 0.19 \pm 0.04\%$; Figure 21.B).

We reanalysed the difference in ghrelin levels between conditions, now by controlling for the glucose levels. For this, we compared a Bayesian linear model with glucose as a predictor for ghrelin to a model with glucose and the condition as predictive values. When controlled for glucose, there was strong evidence in favour of a difference in ghrelin levels between conditions in study 1 ($BF = 36.81 \pm 0.81\%$). However, there was anecdotal evidence against a difference in ghrelin between conditions in study 2 ($BF = 0.49 \pm 1.84\%$).

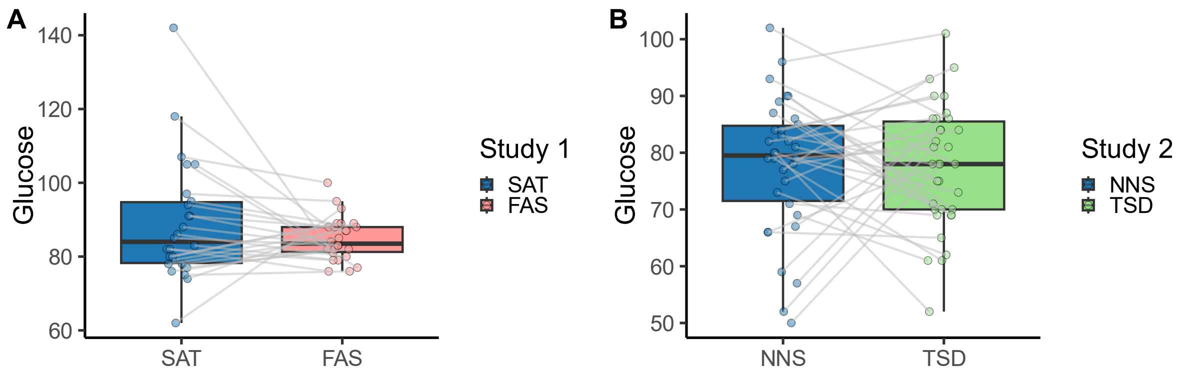


**Figure 21.** Glucose Levels. A) Glucose did not differ between the sated ($Mdn_{SAT} = 80$) and fasted condition (red; $Mdn_{FAS} = 76$) in study 1. B) There were no changes in glucose levels between one night of normal sleep (blue; $Mdn_{NNS} = 79.5$) or one night of total sleep deprivation (green; $Mdn_{TSD} = 78$) in study 2.
